# Dynamic remodeling of ribosomes and endoplasmic reticulum in axon terminals of motoneurons

**DOI:** 10.1101/2021.10.12.463928

**Authors:** Chunchu Deng, Mehri Moradi, Sebastian Reinhard, Changhe Ji, Sibylle Jablonka, Luisa Hennlein, Patrick Lüningschrör, Sören Doose, Markus Sauer, Michael Sendtner

## Abstract

In neurons, endoplasmic reticulum forms a highly dynamic network that enters axons and presynaptic terminals and plays a central role in Ca^2+^ homeostasis and synapse maintenance. However, the underlying mechanisms involved in regulation of its dynamic remodeling as well as its function in axon development and presynaptic differentiation remain elusive. Here, we used high resolution microscopy and live cell imaging to investigate rapid movements of endoplasmic reticulum and ribosomes in axons of cultured motoneurons after stimulation with Brain-derived neurotrophic factor. Our results indicate that the endoplasmic reticulum extends into axonal growth cone filopodia where its integrity and dynamic remodeling are regulated mainly by actin and its motor protein myosin VI. Additionally, we found that in axonal growth cones, ribosomes assemble into 80S subunits within seconds and associate with ER in response to extracellular stimuli which describes a novel function of axonal ER in dynamic regulation of local translation.

**Summary statement:** In axonal growth cones of motoneurons, dynamic remodeling of ER is coordinated through drebrin A-mediated actin and microtubule crosstalk and contributes to local translation as response to BDNF stimulation.

## Introduction

In neurons, the endoplasmic reticulum (ER) provides a luminal space throughout the cytoplasm, which extends into dendrites and axons (Terasaki, Slater et al. 1994, Wu, Whiteus et al. 2017). Within presynaptic terminals, the ER forms a network with predominant tubular appearance close to the active zone, which is highly dynamic and undergoes constant movement and reorganization and regularly forms contact sites with the plasma membrane (Wu, Whiteus et al. 2017, Cohen, Valm et al. 2018). The dynamic movements of ER are regulated by the cytoskeleton and motor proteins. Live cell fluorescence microscopy studies have shown that microtubules regulate ER movements in animal cells (Waterman-Storer and Salmon 1998, Lu, Ladinsky et al. 2009, Wozniak, Bola et al. 2009, Friedman, Webster et al. 2010), whereas in plants (Griffing 2010) and budding yeast cells (Prinz, Grzyb et al. 2000, Du, Walker et al. 2006) ER movements require actin. Similarly, in neurons, microtubules provide a structural backbone for gross dendritic and axonal ER movements (Farias, Freal et al. 2019). Other studies revealed a specific role of myosin Va in ER transport along actin filaments into dendritic spines, demonstrating that ER import into these subcellular compartments might be actin dependent (Wagner, Brenowitz et al. 2011). Whether similar actin dependent mechanisms also determine the movement of axonal ER remains unclear. In this study, we show that in axons, the ER appears associated with actin filaments, especially in growth cone filopodia where actin filaments are highly enriched. The importance of the axonal ER dynamic regulation is highlighted by mutations in ER-shaping or ER-receptor proteins which impair ER remodeling and associate with neurodegenerative diseases such as amyotrophic lateral sclerosis (ALS) (Teuling, Ahmed et al. 2007) or spastic paraplegia (HSP) (Blackstone 2012, Öztürk, O’Kane et al. 2020).

In neurons, rough cisternal ER (RER), which is the major site for protein synthesis, folding, processing and secretion appears mainly restricted to the somatodendritic compartment (Horton and Ehlers 2003, Shibata, Shemesh et al. 2010, West, Zurek et al. 2011, Puhka, Joensuu et al. 2012, Lee, Cathey et al. 2020). Axons are traditionally considered to be devoid of RER, as demonstrated by ultrastructure EM and thought to exhibit only smooth ER (Tsukita and Ishikawa 1976, Krijnse-Locker, Parton et al. 1995). The function of such axonal ER has been suggested to be limited to lipid metabolism, Ca^2+^ homeostasis, and to functions in contacting membranous organelles to regulate their biogenesis and maintenance (Tsukita and Ishikawa 1976, Wu, Whiteus et al. 2017, Farias, Freal et al. 2019, Lee, Cathey et al. 2020). Nevertheless, despite emerging evidence of intra-axonal translation of mRNAs encoding membrane and secreted proteins in isolated neurons (Merianda, Lin et al. 2009), there is no direct evidence for the existence of a rough ER in axons that could process such locally synthesized proteins for integration into the axoplasmic membrane and secretion.

Here, we investigated the interaction of ribosomes with ER in the axonal growth cones of cultured motoneurons and found that ribosomes undergo rapid changes in distribution and structure in response to extracellular cues. We used culture conditions with laminin-2 which promote differentiation of presynaptic structures in axon terminals (Jablonka, Beck et al. 2007). In such differentiated growth cones with presynaptic structures, ribosomes relocate to the ER where they could accomplish local translation of membrane-associated and secreted proteins including TrkB and N-type Ca^2+^ channels. Furthermore, we unraveled the underlying mechanisms for regulation of ER dynamic movements in axon terminals and showed that fast dynamic elongation of ER into axonal filopodia is regulated mainly by actin and its motor protein myosin VI. On the other hand, slow ER movements in the growth cone core depend on a coordinated actin and microtubule cytoskeleton that requires the crosstalk activity of drebrin A. Thus, we provide evidence of a novel function of axonal ER in local protein synthesis of transmembrane proteins such as the α-1β subunit of presynaptic N-type Ca^2+^ channels in addition to its previously described roles.

## Results

### ER dynamics are regulated by a coordinated actin/microtubule cytoskeleton in growth cones of cultured motoneurons

In axons, membrane-bound organelles representing ER often colocalize with microtubules and hardly extend beyond microtubules (Dailey and Bridgman 1989, Farias, Freal et al. 2019). Nevertheless, growth cone filopodia that contain differentiated presynaptic active zone structures (Jablonka, Beck et al. 2007) are enriched in actin filaments and mostly lack microtubules (Geraldo and Gordon-Weeks 2009). Thus, we wondered whether ER in axonal growth cones could extend into these actin-rich filopodia and associate with actin filaments in addition to microtubules. In order to visualize the ER in growth cones, we transduced primary cultured motoneurons with a lentivirus expressing mCherry-KDEL, a well-studied ER marker (Guo, Li et al. 2018). Neurons were then fixed and immunostained to visualize F-actin and the microtubule cytoskeleton using phalloidin and α-tubulin staining, respectively. The association of ER with actin and tubulin in axon terminals was then assessed using Structured Illumination Microscopy (SIM) (Fig. S1). Interestingly, ER was not only detected in the core of growth cones but also in filopodia (Fig. S1A). 3D reconstruction and line scan analysis showed that ER colocalizes with both F-actin and microtubules in the core, while in filopodia, ER overlaps mostly with F-actin, as expected since microtubules are less abundant in filopodia (Fig. S1B,C). To rule out colocalization by chance, we rotated the ER channel by 90 degree and could detect only a small overlap between the ER and either F-actin or α-Tubulin (Fig. S1D). Next, we performed live cell imaging over a period of 8 min with 2 sec intervals to visualize ER movements in the axonal growth cone. We transduced motoneurons with mCherry-KDEL lentivirus and co-transduced with a lentivirus expressing a cell volume marker (GFP) for simultaneous visualization of the plasma membrane movements (Fig. 1A; movie 1). Notably, ER entered only a fraction of highly dynamic filopodia in axonal growth cones during the time of observation, indicating that ER movement is only partially coupled to movements of the plasma membrane (Fig. 1B). To confirm the colocalization of the ER with F-actin in filopodia, we monitored the dynamics of ER/actin co-movements in filopodia by transducing motoneurons with a GFP-actin-IRES-mCherry-KDEL lentivirus. This lentiviral construct expresses both GFP-actin and mCherry-ER and thus allows visualization of actin and ER movements in the same filopodia simultaneously (Fig. 1C,D). Two color live cell imaging demonstrated that in filopodia, tubular ER extends and retracts along actin filaments indicating that ER interaction with actin defines its rearrangement and remodeling (movie 2). It is of note that in some filopodia, ER collapses while actin filaments remain stable (movie 3). To quantify ER dynamic movements, we implemented an adaption of Image Correlation Spectroscopy (ICS) (Wiseman 2015), which measures the dynamics as intensity movements (µm/sec) (Fig. 1E). ICS-analysis revealed distinct motilities for actin and ER in the same filopodia which excludes the possibility that ER extension/retraction events are forced by actin movements (Fig. 1F). Based on these observations, we hypothesized that ER extension into filopodia might be actin and not microtubule dependent. To address this, we transduced motoneurons with mCherry-KDEL and disrupted F-actin or microtubules using depolymerizing drugs cytochalasin D (CytoD) or nocodazole, respectively. Motoneurons were treated with 1 µg/ml CytoD for 30 min or 10 µM nocodazole for 2 h and ER movements were then evaluated in the growth cone by live cell imaging (Fig. 2A,B; movie 4-9). ICS-analysis did not detect any movements in fixed cells, indicating that the error of ICS-analysis is very low (Fig. 2C). In addition to ICS-analysis, we used multiple kymographs to measure the frequency and distance moved by ER in the axonal growth cones manually. Intriguingly, we found that the dynamic movement of ER in filopodia is significantly higher than that in the core (Fig. 2C). Besides, the velocity of dynamic movements was higher, as represented in Fig. 2D, and the distance moved by ER in filopodia was greater compared to growth cone core, correspondingly (Fig. 2E). Moreover, we found that upon CytoD treatment, 80% of neurons failed to show ER movements in filopodia, while only 40% failed to show ER movements in the core (Fig. 2F). Treatment with nocodazole resulted in 50% failure in filopodia and 40% failure in core ER movements (Fig. 2F). ICS-analysis revealed that disruption of either actin or microtubule dynamics reduces ER movements significantly in both filopodia and core implying that ER dynamic movements depend on a coordinated actin/microtubule cytoskeleton (Fig. 2G,H). Nevertheless, analyzing the frequency of ER movements in filopodia demonstrated that disruption of actin but not microtubules reduced the frequency of ER movements (Fig. 2G). In contrast to filopodia, the frequency of ER movements in core was markedly reduced upon disruption of either actin or microtubule cytoskeleton which is in agreement with the above ICS-analysis (Fig. 2H). This finding was further confirmed by a treatment with both CytoD and nocodazole which severely impaired the ER dynamics in both filopodia and core, as nearly no ER movements were detectable under this condition (Fig. 2F-H; Fig. S2A).

**Figure 1.**
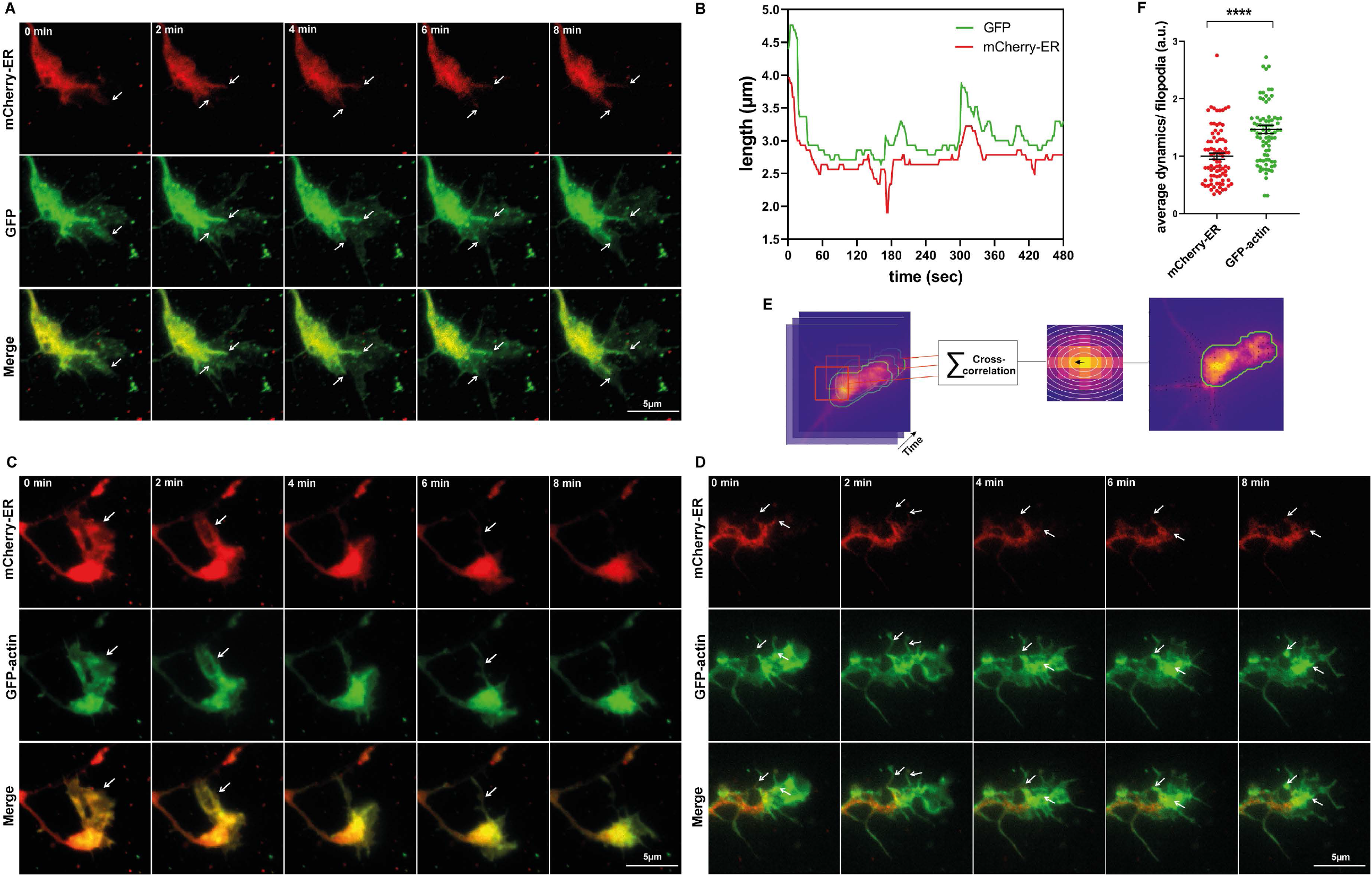
ER moves along actin filaments in growth cone filopodia. (A) Representative time-lapse images of motoneurons transduced with lentiviruses expressing cell volume marker GFP and mCherry-KDEL. Arrows indicate ER entry into filopodia that are labeled with GFP. (B) Graph shows representative growth and retraction events of ER and plasma membrane (GFP) in a single filopodia over time. (C and D) Representative time-lapse images of motoneurons transduced with a lentivirus co-expressing GFP-actin and mCherry-KDEL. (C) Arrows indicate co-movements of mCherry-ER and GFP-actin in some filopodia. (D) Arrows indicate that in some filopodia only ER but not F-actin retracts. (E) Diagram shows workflow of ICS python implementation. (F) Graph shows average dynamics of ER and F-actin movements per filopodia. Dynamic movements of F-actin are significantly higher than those of ER in the same filopodia (****, P<0.0001; n=83 filopodia in 3 independent experiments). Graph is shown in scatter dot plot with mean±SEM. Statistical analyses: two-tailed Mann Whitney test.

**Figure 2.**
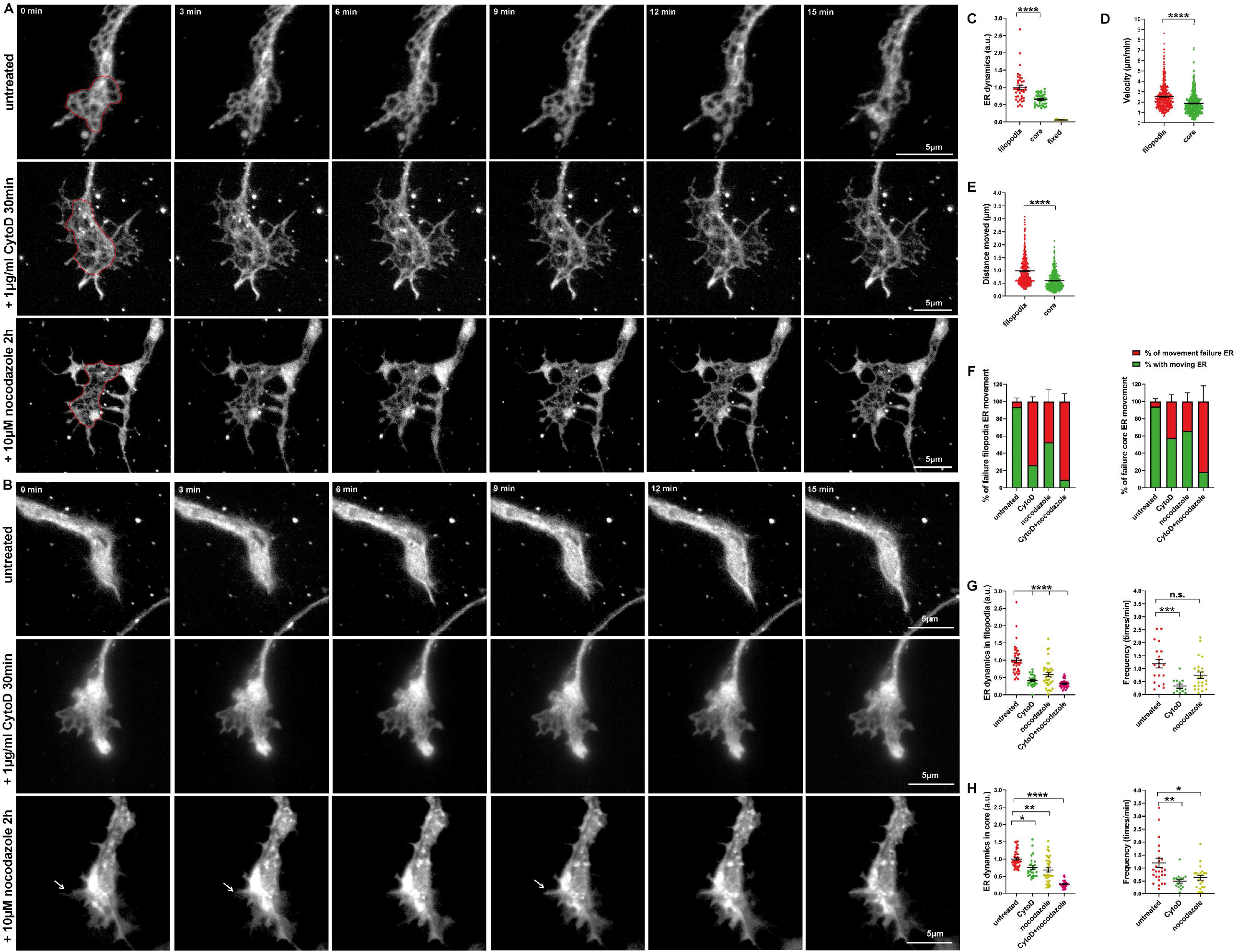
ER dynamic movements are regulated by actin and microtubules in growth cone core and filopodia. (A and B) Representative time-lapse images of motoneurons expressing mCherry-ER in the growth cone core (A) and filopodia (B). Red circles mark growth cone core and arrows show ER extension into filopodia. (C) Graph shows ICS-analysis of ER movements in filopodia versus core. ER dynamic movements are higher in filopodia versus core (****, P<0.0001; n=40 cells from 6 independent experiments). (D and E) Kymograph-analysis of velocity of ER movements (D) and distance moved by ER (E) in filopodia versus core (****, P<0.0001; n=25-29 cells from 3 independent experiments). (F) Average percentage of neurons failing to show ER movements in filopodia or core upon treatments (n=32-55 cells from 3 independent experiments). (G) Average ER movements (left panel: ****, P<0.0001; n=27-40 cells from 4-5 independent experiments) and frequency of ER movements (right panel) in filopodia per growth cone. In filopodia, frequency of ER movements is decreased upon CytoD (***, P=0.0004; n=12-20 cells), but not nocodazole treatment (n.s., P=0.1079; n=20-23 cells from 4-5 independent experiments). CytoD+nocodazole treatment decreases ER movements in filopodia (****, P<0.0001; n=28-40 cells from 3 independent experiments). (H) Average ER movements (left panel: *, P=0.0353; **, P=0.0012; n=27-40 cells from 4-5 independent experiments) and frequency of ER movements (right panel: **, P=0.0016; *, P=0.0271; n=15-27 cells from 4-6 independent experiments) in core per growth cone. CytoD+nocodazole treatment decreases core ER movements (****, P<0.0001; n=28-40 cells from 3 independent experiments). Graphs are shown in bar diagrams or scatter dot plot with mean±SEM. Statistical analyses: one-way ANOVA with Dunn’s post-test in G and H, and two-tailed Mann Whitney test in C-E.

Collectively, these data provide evidence of a highly dynamic ER in axonal growth cones of developing motoneurons. The dynamic movements of ER in axonal growth cones could be classified into fast movements in filopodia and slower movements in the core. Fast ER remodeling in filopodia is regulated particularly by the actin cytoskeleton, whereas slow ER rearrangements in the core require a microtubule and actin crosstalk.

### Myosin VI particularly tethers the ER into axonal filopodia

Next, we asked which myosin motor protein is involved in actin-mediated ER tethering into axonal growth cone filopodia. A previous study showed that myosin Va translocates the ER into dendritic spines of Purkinje neurons (Wagner, Brenowitz et al. 2011). To examine the possible role of different myosin isoforms, we treated motoneurons with pharmacological inhibitors of myosin II ((-)-blebbistatin), V (MyoVin-1) and VI (2,4,6-triiodophenol) and analyzed the ER movements in growth cones by live cell imaging (Fig. 3A). First, we tested the toxicity of these inhibitors via survival assay using concentrations ranging between 1 µM and 100 µM and tested incubation times of 30 min up to 12 h. Treatments with 5 µM (-)-blebbistatin, 30 µM MyoVin-1 and 1 µM 2,4,6-triiodophenol for 15 min seemed not to affect the neuronal viability and were therefore chosen for this experiment (Fig. S3A,B,C). Pharmacological inhibition of myosin V and especially myosin VI markedly reduced the dynamic movements of ER in both core and filopodia of growth cones, whereas inhibition of myosin II was much less effective (Fig. 3B,C; movie 10-12). These data indicate that in motoneurons, particularly myosin VI drives ER movements in axonal growth cones.

**Figure 3.**
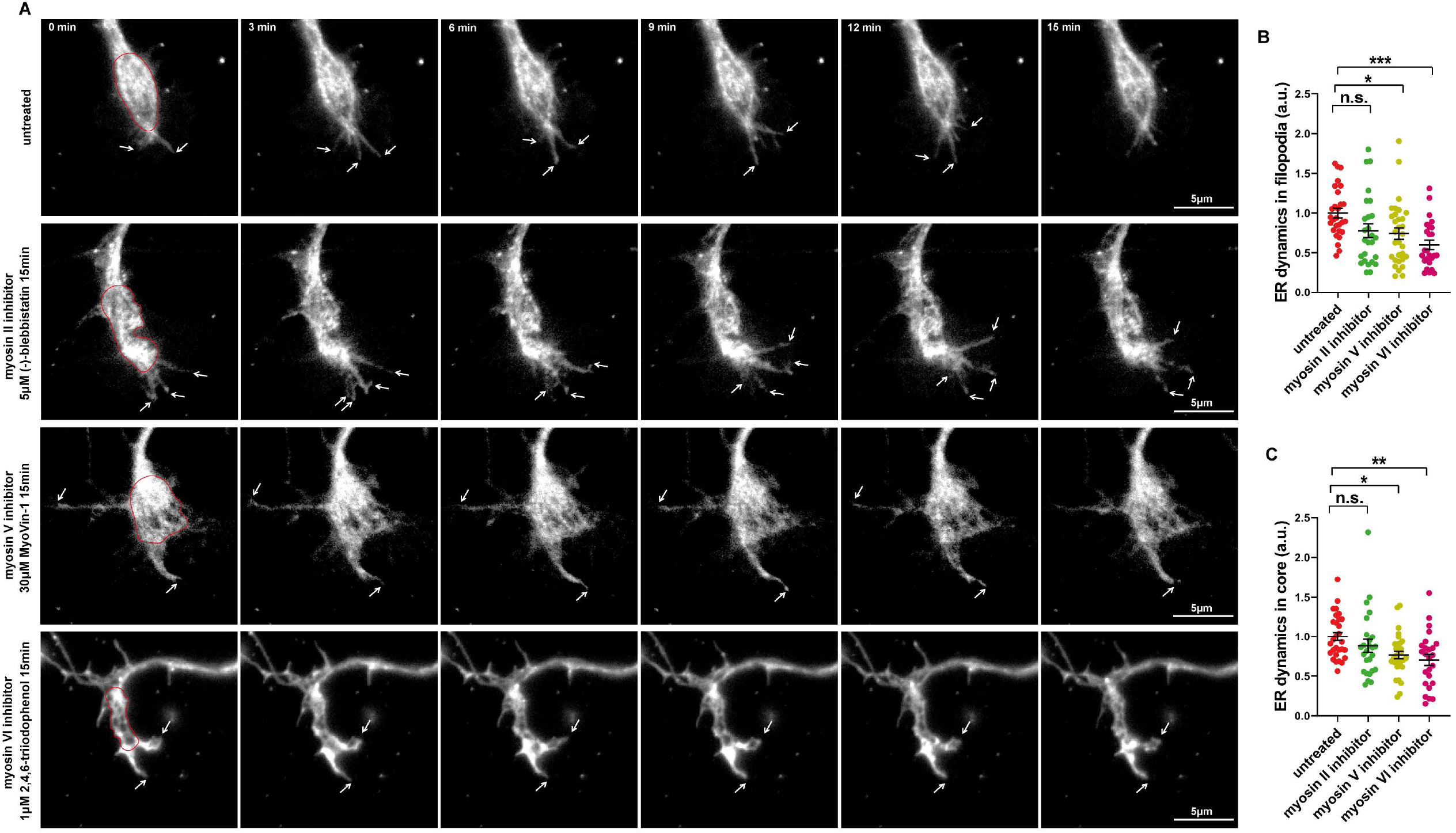
ER entry into axonal growth cone filopodia depends on Myosin VI. (A) Representative time-lapse images of ER in growth cones of motoneurons treated with myosin II, V or VI inhibitors and the untreated control. Red circles mark the ER in growth cone cores and white arrows indicate the movements of filopodia ER. (B) Quantification of ER dynamics in mCherry-KDEL transduced motoneurons reveals significant reduction in ER movements in axonal filopodia upon inhibition of myosin V (*, P<0.0361) and VI (***, P<0.0005; n=26-31 cells from 4 independent experiments).(C) Graph shows reduced ER dynamic movements in the growth cone core after treatments with myosin V (*, P<0.0339) and VI inhibitors (**, P<0.0099; n=26-31 cells from 4 independent experiments). Graphs are shown in scatter dot plot with mean±SEM. Statistical analyses: one-way ANOVA with Dunn’s post-test.

### Drebrin A regulates axonal ER movements via actin and microtubule crosstalk

Previous studies revealed that drebrin mediates the actin and microtubules crosslink through direct interaction with actin filaments and the microtubule end-binding protein 3 (EB3) (Geraldo, Khanzada et al. 2008, Worth, Daly et al. 2013). This drebrin-depending actin/microtubule crosstalk is required for neuronal migration (Trivedi, Stabley et al. 2017) and neuritogenesis (Geraldo, Khanzada et al. 2008). Thus, we sought to examine whether coordination between actin and microtubules which is required for ER movements in axonal growth cones depends on drebrin. Derbrin has two major isoforms; neuronal specific drebrin A which is highly expressed in the adult brain, and embryonic isoform drebrin E which is expressed also in non-neuronal cells (Shirao, Hanamura et al. 2017). We found similar expression levels of these two isoforms in our cultured motoneurons from mouse embryos. To examine the role of these isoforms in axonal ER movements, we designed two different lentiviral shRNAs that target either drebrin A (shDrebrin A) or both drebrin A and E (shDrebrin A+E). These lentiviral constructs co-expressed GFP which was used to identify transduced cells. As control, we used the empty shRNA vector backbone expressing only GFP (shCtrl). qRT-PCR assay showed that transduction of motoneurons with shDrebrin A leads to 90% reduction in drebrin A mRNA levels and transduction with shDrebrin A+E results in 90% reduction in drebrin A and 94% reduction in drebrin E mRNA levels (Fig. 4A). Next, we co-transduced motoneurons with mCherry-ER and either shDrebrin A or shDrebrin A+E, or shCtrl and analyzed the ER dynamic movements in axonal growth cones by live cell imaging (Fig. 4B; movie 13-14). We found that knockdown of drebrin A decreases ER dynamic movements by 40% in filopodia (Fig. 4C) and 30% in the growth cone core (Fig. 4D). As illustrated in Fig. 4C,D, additional knockdown of drebrin E did not further decrease the ER movements, indicating that drebrin A but not drebrin E is relevant for coordinating actin and microtubule functions involved in ER movements in the axonal growth cone.

**Figure 4.**
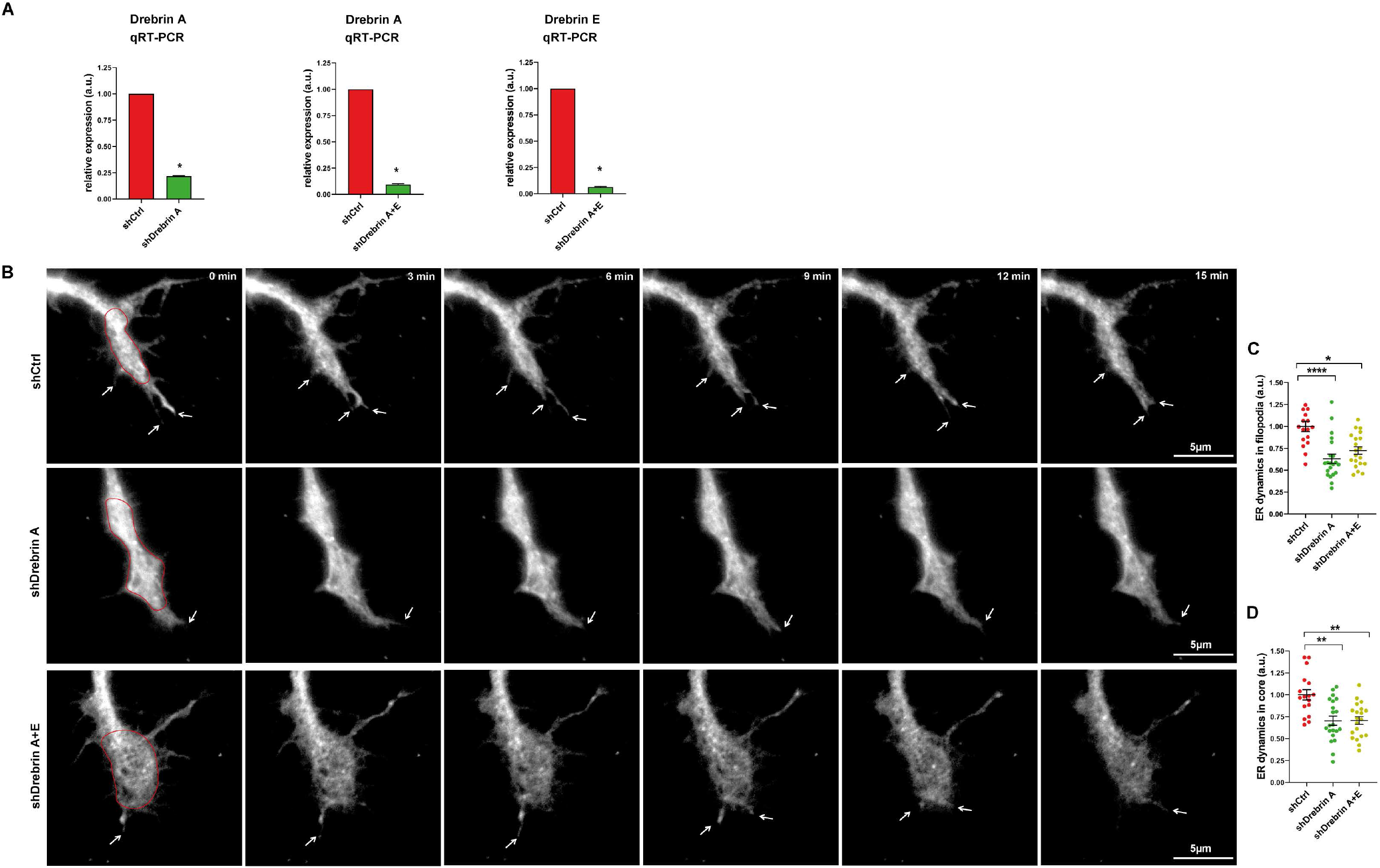
Drebrin A mediates actin and microtubule coordinated ER movements in axonal growth cones. (A) Graphs show relative mRNA expression of drebrin A and E in shDrebrin A and shDrebrin A+E-transduced motoneurons. shRNA-mediated knockdown of drebrin A results in 80% reduction in drebrin A mRNA levels (left panel: *, P<0.05 from 3 independent experiments). shRNA targeting both drebrin A and E (shDrebrin A+E) causes 90% reduction in drebrin A (middle panel: *, P<0.05) and 94% reduction in drebrin E mRNA levels (right panel: *, P<0.05; from 3 independent experiments). (B) Representative time-lapse images of ER in motoneuron growth cones transduced with shDrebrin A, shDrebrin A+E and shCtrl lentiviruses. ER in the growth cone core is marked by red circles and ER movements in filopodia are indicated by white arrows. (C) Graph shows dynamic of ER movements in axonal growth cone filopodia. Knockdown of drebrin A significantly reduces the ER movements in filopodia (****, P<0.0001). Knockdown of drebrin E in addition to drebrin A does not further reduce the ER movements in filopodia (*, P=0.0133; n=17-21 cells from 3 independent experiments). (D) ER movements in the growth cone core are depicted in the graph. Knockdown of drebrin A causes significant reduction in ER movements in the growth cone core (**, P=0.0026), while additional knockdown of drebrin E does not intensify this effect (**, P=0.0019; n=17-21 cells from 3 independent experiments). Graphs are shown in bar diagrams or scatter dot plot with mean±SEM. Statistical analyses: one-tailed Mann Whitney test in A and one-way ANOVA with Dunn’s post-test in C and D.

### Extracellular stimulation triggers ribosome activation and initiates local translation in growth cones on a time scale of seconds

Deep RNA-seq combined with sensitive fluorescence in situ hybridization (FISH) approaches have identified numerous mRNAs (Briese, Saal et al. 2016, Holt, Martin et al. 2019) within distal axons which are locally translated (Moradi, Sivadasan et al. 2017, Terenzio, Koley et al. 2018, Biever, Glock et al. 2020). Locally synthesized proteins are necessary for neural-circuit development, survival and plasticity (Fernandopulle, Lippincott-Schwartz et al. 2021). In developing axons, local translation mediates an essential response to guidance cues required for pathfinding (Leung, van Horck et al. 2006). Brain-derived neurotrophic factor (BDNF) and its receptor TrkB play a vital role in modulation of local translation (Santos, Comprido et al. 2010) and axonal cytoskeleton remodeling (Sasaki, Welshhans et al. 2010, Rathod, Havlicek et al. 2012) in motoneurons. We sought to scrutinize the dynamics of ribosome activation in response to BDNF stimulation in the growth cone of motoneurons. Following BDNF binding, TrkB undergoes autophosphorylation and activates MAPK as well as PI3K-AKT signaling pathways (Huang and Reichardt 2003, Hua, Gu et al. 2016), leading to activation of ribosomes and induction of local translation (Leal, Comprido et al. 2014). In order to define the precise kinetics of TrkB activation, we applied BDNF to motoneurons with a short pulse of 10 sec, 1 min and 10 min, washed it out and immunostained neurons against TrkB and pTrkB. The specificity of TrkB and pTrkB antibodies was confirmed by immunostaining using TrkB knockout mice (Fig. S4A,B). We could detect phosphorylated TrkB upon 10 sec BDNF pulse in the growth cone, as shown by immunofluorescence assay (Fig. 5A,B). Interestingly, the levels of TrkB also elevated in the growth cone after 1 min BDNF exposure (Fig. 5C). In the whole cell lysate, a corresponding elevation of pTrkB immunoreactivity became detectable first at 1 min poststimulation, as shown by Western blot (Fig. 5D) and no increase in total levels of TrkB was detectable despite 10 min stimulation, indicating that this short pulse is insufficient to induce the transcription and translation of TrkB in the soma (Fig. 5D; Fig. S4C). The rapid increase in TrkB immunoreactivity within less than 1 min in the growth cone of motoneurons could be explained by release of this receptor from intracellular stores or changes in the receptor conformation, which favors antibody binding. The increase in TrkB immunoreactivity in growth cones at a later time point of 10 min poststimulation (Fig. 5C) could be explained by enhanced transport of TrkB from distal axons into growth cones and/or rapid neosynthesis in growth cones since TrkB mRNAs were detected in axons by qRT-PCR assay (Fig. S4D). Treatment of neurons with either 100 ng/ml anisomycin for 1 h, which inhibits the translation, or 10 µM nocodazole for 2 h, which blocks the microtubule-based transport, prevents the increase in TrkB signal in growth cones (Fig. 5E,F). These data indicate that BDNF stimulation triggers redistribution of TrkB resulting in an increase in TrkB total immunofluorescence within 1 min, and also induces its local production within 10 min stimulation.

**Figure 5.**
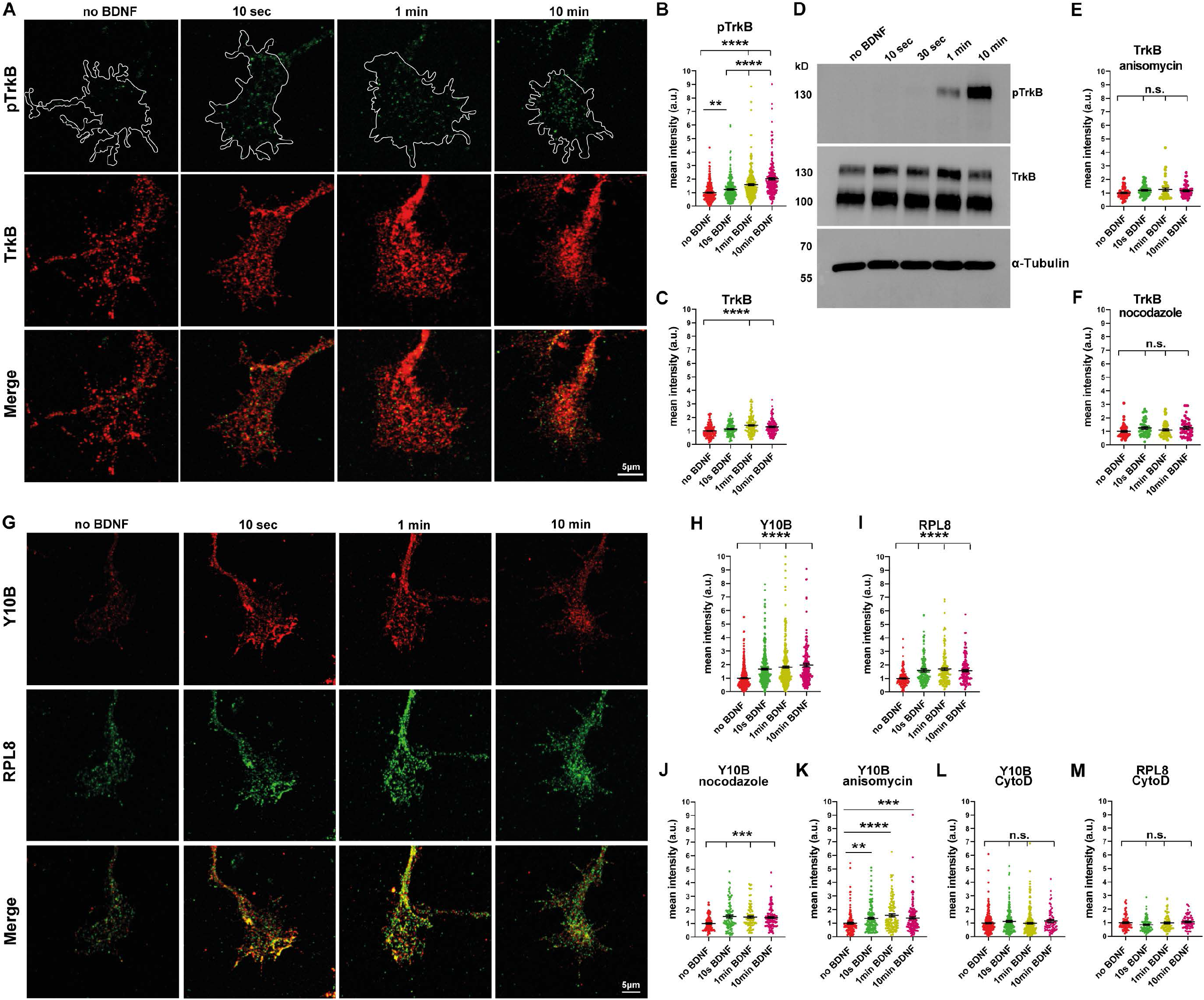
BDNF-induced TrkB activation triggers redistribution of ribosomes in growth cones. (A) Representative images of growth cones of BDNF-stimulated motoneurons stained against pTrkB and TrkB. (B and C) Mean intensities of pTrkB increase at 10 sec (**, P=0.0063; n=265-267 cells) and mean intensities of pTrkB and TrkB increase at 1 min (****, P<0.0001; n=123-274 cells) and 10 min poststimulation (****, P<0.0001; n=120-266 cells from 8 and 4 independent experiments). (D) Representative immunoblot of total lysates from cultured motoneurons probed against pTrkB, TrkB and α-Tubulin. (E and F) Total levels of TrkB in stimulated growth cones treated with nocodazole (n.s., P≥0.1526; n=45-46 cells) or anisomycin (n.s., P≥0.0893; n=47-54 cells from 2 independent experiments). (G) Representative images of stimulated growth cones stained against Y10B and RPL8. (H and I) Mean intensities of Y10B and RPL8 increase at 10 sec (****, P<0.0001; n=138-366 cells from 9 and 4 independent experiments), 1 min (****, P<0.0001; n=138-366 cells from 9 and 4 independent experiments) and 10 min poststimulation (****, P<0.0001; n=138-210 cells from 6 and 4 independent experiments). (J and K) Y10B immunoreactivity increases despite nocodazole (***, P≤0.0004; n=93-102 cells from 3 independent experiments) or anisomycin treatment (**, P=0.0012; ****, P<0.0001; ***, P=0.0008; n=121-141 cells from 5 independent experiments). (L and M) Y10B (n.s., P≥0.5207; n=68-252 cells from 3-6 independent experiments) and RPL8 immunoreactivities (n.s., P≥0.2256; n=68-87 cells from 3 independent experiments) do not increase with CytoD treatment. All data are normalized to the average intensities of no BDNF group. Graphs are shown in scatter dot plot with mean±SEM. Statistical analyses: one-way ANOVA with Dunn’s post-test.

In developing axons, the rapid response to extracellular cues is assured by tight regulation of spatiotemporal changes in mRNA translation (Willis, van Niekerk et al. 2007, Yoon, Zivraj et al. 2009). Thus, the kinetics of induction of ribosomal changes that trigger translation in growth cones should differ from those in the soma. To study the dynamics of ribosome remodeling, we used Y10B as a general marker for ribosomes and RPL8 as a marker for the 60S ribosomal subunit in order to study the dynamics of movement of these different subunits. We found that induction of BDNF/TrkB signaling leads to a rapid change in the distribution of these ribosomal markers within the growth cone. The immunoreactivities of ribosomal markers Y10B and RPL8 were rapidly altered and appeared increased after 10 sec BDNF pulse (Fig. 5G-I). Interestingly, short pulses of 10 sec and 1 min were not sufficient to induce such changes in ribosome distribution in the soma as shown in Fig. S4E, albeit TrkB phosphorylation was detectable also in this subcellular compartment after 10 sec stimulation (Fig. S4F). This kinetic distinction suggests that in the growth cone, BDNF/TrkB signaling and downstream mechanisms for modulating ribosome distribution and possibly also activation are different from cell bodies. Increased immunoreactivity for ribosomal subunits in the growth cone within 10 sec BDNF stimulation is striking. Rapid transport of ribosomal subunits from axonal sites to the growth cone appears unlikely since the fastest measured microtubule-dependent axonal transport at a speed of 4.6 μm/sec cannot provide ribosomes from the axon shaft into growth cones within such a short time (Maday, Twelvetrees et al. 2014, Wortman, Shrestha et al. 2014). To exclude this possibility, we treated neurons with nocodazole to disrupt microtubule-based axonal transport and found that disruption of axonal transport did not affect BDNF-induced augmentation of ribosomal subunits as illustrated in Fig. 5J. Similarly, inhibition of translation by anisomycin treatment results in a significant increase in ribosome immunoreactivity, indicating that this increase does not depend on de novo synthesis of ribosomal proteins within the growth cone (Fig. 5K). Strikingly, treatment with 1 µg/ml CytoD for 30 min completely prevents the ribosomal response to BDNF stimulation, as no increase in the signal intensities of Y10B and RPL8 were detectable upon either stimulation interval (Fig. 5L,M). Based on these observations, we propose that elevated immunoreactivity of ribosomes within 10 sec stimulation might be due to actin dependent conformational and distribution changes in ribosomal subunits which occurs upon assembly of 80S subunits and thus alters accessibility of corresponding epitopes for detection with Y10B and RPL8 antibodies. To investigate this hypothesis, we stained motoneurons with antibodies against RPL24 of the 60S and RPS6 of the 40S subunits (Fig. 6A). Upon assembly of 80S ribosomes, the C-terminus of RPL24 interacts with RPS6, thereby forming a bridge between the two subunits (Ben-Shem, Garreau de Loubresse et al. 2011). We employed SIM with a resolution of ∼120 nm to examine the interaction of RPL24/RPS6 in growth cones. As depicted in Fig. 6B, within 10 sec of stimulation the co-clusters of RPL24/RPS6 increased by 3-fold compared to no stimulation condition. Similarly, colocalization of RPL24 and RPS6 increases significantly at 10 sec and 1 min poststimulation (Fig. 6C). Thus, we reasoned that the rapid increase in Y10B immunoreactivity in response to extracellular stimulation involves extremely fast assembly of ribosomes into 80S subunits. Strikingly, 80S subunits seem to disassemble after a long stimulation of 30 min as the number of RPL24/RPS6 co-clusters as well as their colocalization declined. This observation implies a rapid but transient response of ribosomes to extracellular stimuli (Fig. 6B,C). Strikingly, inhibition of actin polymerization via CytoD treatment impeded the formation of RPL24/RPS6 co-clusters (Fig. 6D). Consistent with the data represented in Fig. 5L,M, these data clearly show that ribosome distribution and assembly depend on the actin cytoskeleton. The increased interaction of RPL24 and RPS6 ribosomal proteins and consequently the assembly of 80S ribosomes suggest that this conformational change could correlate with translation initiation. To visualize actively translating ribosomes, we co-stained motoneurons against Y10B as a structural component of ribosomes and the elongation factor eEF2, which associates only with actively translating ribosomes (Fig. 6E). Again, we found a significant increase in the number of Y10B/eEF2 co-clusters within 10 sec of stimulation, indicating that the response of ribosomes in growth cones to activation of TrkB receptors is very fast (Fig. 6F). Colocalization analysis confirmed increased association of eEF2 to Y10B at 10 sec and 1 min stimulations (Fig. 6G). To confirm that the rapid activation of ribosomes depends on BDNF/TrkB signaling, we stained motoneurons from TrkB knockout mice against Y10B and found that in TrkB knockout motoneurons ribosomes fail to respond to 10 sec BDNF stimulation (Fig. S4G,H). Next, we questioned whether this rapid activation of ribosomes in response to extracellular stimuli associates with a temporal change in protein synthesis in axonal growth cones. To examine the dynamics of local translation, we applied a time series after BDNF pulse to motoneurons, fixed them and stained against β-actin (Fig. 7A). Remarkably, we observed a significant increase in β-actin signal intensity within 1 min of stimulation suggesting that either the velocity of protein synthesis is extremely high in growth cones or actin undergoes conformational changes in such a way that this leads to enhanced immunoreactivity (Fig. 7B). To distinguish these two possibilities and to confirm that elevated levels of β-actin rely on protein synthesis and not transport from the axon, we pretreated neurons with anisomycin and nocodazole as described above and applied BDNF pulses for 10 sec to 10 min. Inhibition of protein synthesis prevented the increase of β-actin protein levels, while disruption of microtubule-dependent axonal transport had no influence (Fig. 7C,D). Next, we considered the dynamic local translation of transmembrane proteins such as N-type Ca^2+^ channels (Cav2.2), whose mRNAs localizes to distal axons in cultured motoneurons (Briese, Saal et al. 2016). Using an antibody against the α-1β subunit of N-type Ca^2+^ channels, we detected increased levels of α-1β subunits within 1 min stimulation in axonal growth cones (Fig. 7E,F). Treatment of the cells with anisomycin diminished this enhancement (Fig. 7G), whereas treatment with nocodazole still resulted in elevated levels of α-1β subunits within 1 min in response to BDNF (Fig. 7H). In addition, we incubated cells with 10 µg/ml puromycin for 10 min to confirm our results (Fig. 7I,J). Similar to β-actin and α-1β subunits of the N-type Ca^2+^ channels, within 1 min of stimulation, synthesis of de novo proteins was accomplished as detected by marked increase in puromycin immunoreactivity, supporting the observation of an extremely fast rate of local translation in growth cones (Fig. 7K). In contrast, 1 min likewise 10 min stimulations did not trigger translation as measured by puromycin intensity in the soma under the same conditions, pointing again towards different translation kinetics in these distinct compartments (Fig. S4I). In line with that, inhibiting translation but not axonal transport prohibited elevated protein synthesis in response to extracellular stimuli as demonstrated by the puromycin assay (Fig. 7L,M). Taken together, these data indicate that in axon terminals, BDNF/TrkB signaling activates the translational machinery within seconds and thus regulates local protein synthesis at an extraordinary high speed.

**Figure 6.**
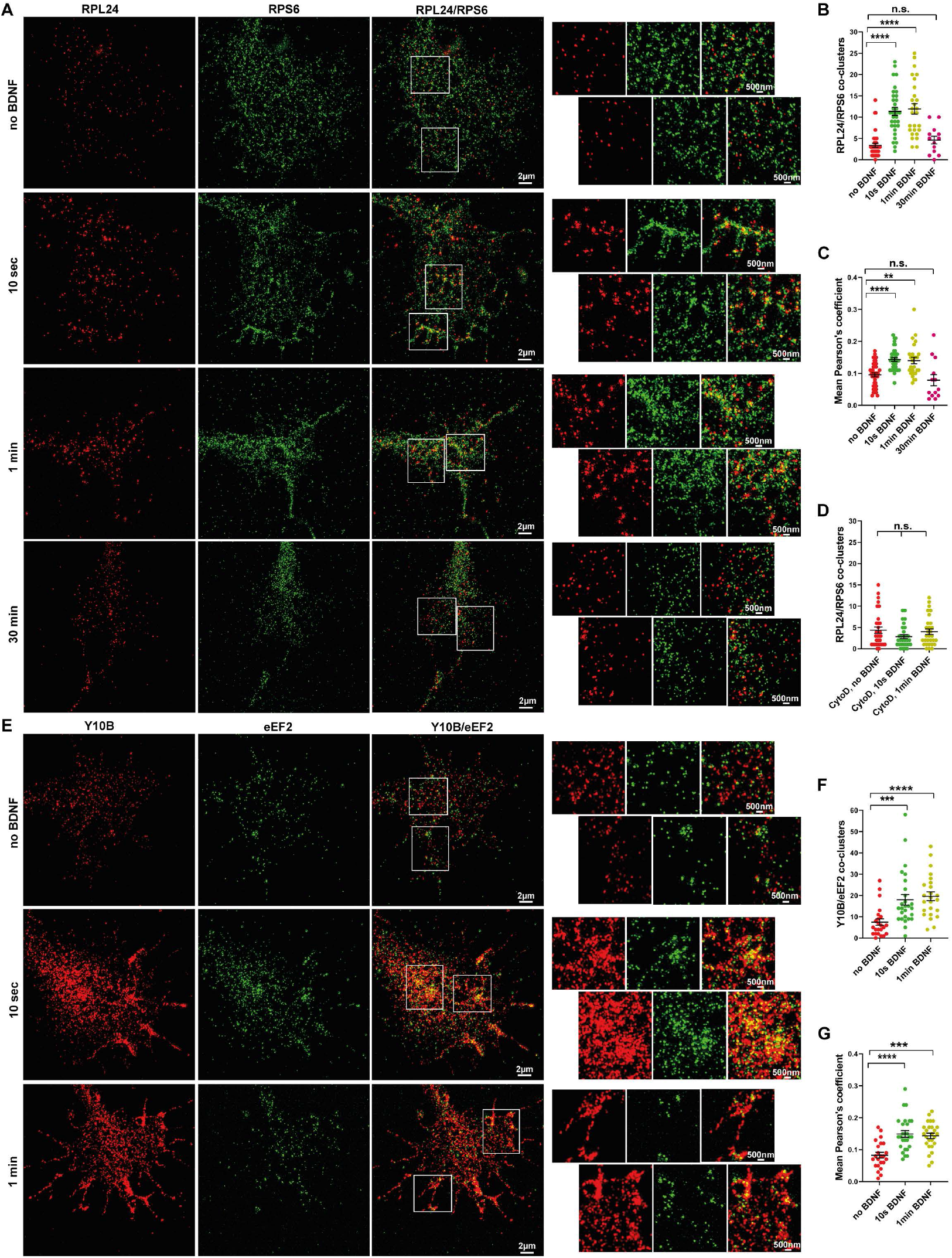
BDNF stimulation induces ribosomal assembly and initiates translation in growth cones of motoneurons. (A) Representative SIM images of growth cones of BDNF-stimulated WT motoneurons stained against RPL24 and RPS6. White squares indicate magnification of ROIs in growth cones. (B) Quantification of RPL24/RPS6 co-clusters that are representative of fully assembled 80S subunits shows that BDNF stimulation induces formation of RPL24/RPS6 co-clusters at 10 sec and 1 min (****, P<0.0001; n=27-33 cells from 3 independent experiments) which dissociate again after a long stimulation of 30 min (n.s., P>0.99). (C) Pearson’s correlation coefficient analysis shows increased colocalization of RPL24 and RPS6 at 10 sec and 1 min (****, P<0.0001; **, P=0.0079; n=26-36 cells from 3 independent experiments) but not 30 min (n.s., P>0.99) poststimulation. (D) Quantification shows that CytoD treatment affects the BDNF-induced formation of RPL24/RPS6 co-clusters (n.s., P=0.616; n=30-32 cells from 3 independent experiments). (E) Representative SIM images of growth cones of BDNF-stimulated motoneurons stained against Y10B and eEF2. White squares indicate magnification of ROIs in growth cones. BDNF stimulation induces formation of Y10B/eEF2 co-clusters at 10 sec as well as 1 min. (F) Quantification of Y10B/eEF2 co-clusters that are representative of ribosomes in the elongation phase of translation (***, P=0.0007; ****, P<0.0001; n=23-27 cells from 3 independent experiments). (G) Mean Pearson’s correlation coefficient shows increased colocalization of Y10B and eEF2 at 10 sec and 1 min poststimulation (****, P<0.0001; ***, P=0.0001; n=23-27 cells from 3 independent experiments). Graphs are shown in scatter dot plot with mean±SEM. Statistical analyses: one-way ANOVA with Dunn’s post-test.

**Figure 7.**
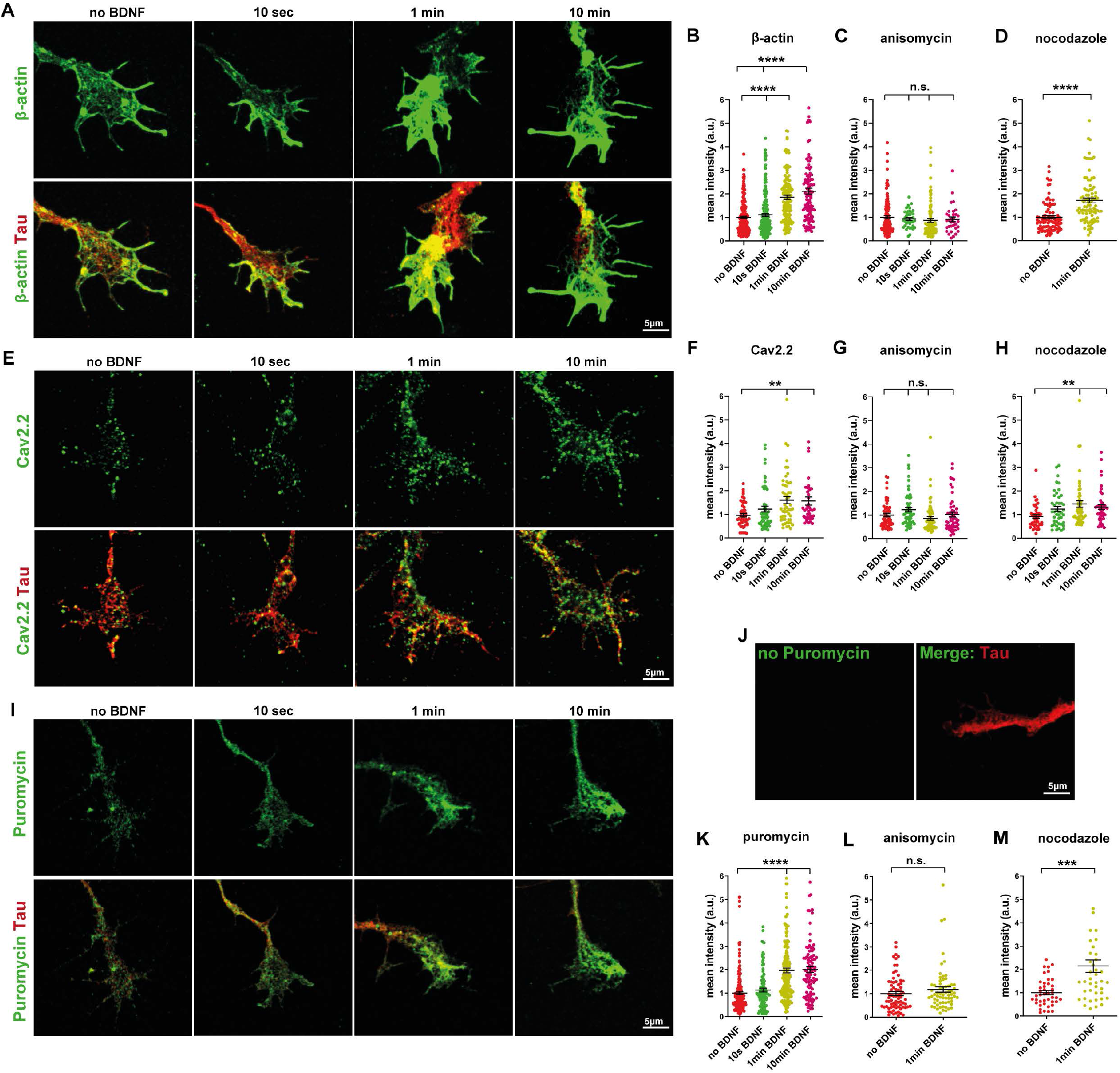
Rapid effect of BDNF stimulation on protein synthesis in motoneuron growth cones. (A) Representative confocal images of growth cones of BDNF-stimulated motoneurons stained against β-actin (green) and Tau (red). (B) Graph shows mean intensities of β-actin in growth cones. A substantial enhancement in mean intensity of β-actin immunoreactivity is first detected after 1 min or 10 min stimulation (****, P<0.0001; n=114-202 cells from 4 independent experiments). (C) Anisomycin-treated cells do not show increased levels of β-actin in the growth cone in response to BDNF stimulation (n.s., P≥0.2903; n=33-127 cells from 2-3 independent experiments). (D) 1 min BDNF stimulation leads to an increase in β-actin protein levels in nocodazole-treated cells (****, P<0.0001; n=91-95 cells from 3 independent experiments). (E) Representative confocal images of growth cones of BDNF-stimulated motoneurons stained against Cav2.2 (green) and Tau (red). (F) Mean intensities of Cav2.2 increase significantly at 1 min (**, P<0.0057) as well as 10 min (**, P<0.0085; n=50-54 cells from 3 independent experiments) poststimulation. (G) Anisomycin treatment abolishes the effect of BDNF on Cav2.2 levels in the growth cone (n.s., P=0.26; n=53-67 cells from 3 independent experiments). (H) Mean intensities of Cav2.2 increase significantly after 1 min (**, P<0.0035) and 10 min (**, P<0.0099; n=43-50 cells from 3 independent experiments) BDNF pulses in growth cones of nocodazole-treated cells. (I) Representative confocal images of growth cones of BDNF-stimulated motoneurons stained against puromycin (green) and Tau (red). (J) As control, puromycin was omitted and cells were incubated only with primary and secondary antibodies against puromycin. (K) Puromycin immunoreactivity levels are increased at 1 min and 10 min post-stimulation (****, P<0.0001; n=103-218 cells from 3-6 independent experiments). (L and M) Anisomycin (n.s., P=0.1796; n=65-73 cells from 2 independent experiments) but not nocodazole (***, P=0.0002; n=42 cells from 2 independent experiments) treatment inhibits puromycin immunoreactivity after BDNF stimulation. All data are normalized to the average intensities of no BDNF group. Graphs are shown in scatter dot plot with mean±SEM. Statistical analyses: one-way ANOVA with Dunn’s post-test in B, C, F, G, H and K, and by two-tailed Mann Whitney test in D, L and M.

### Rough ER is present in the axonal growth cone and contributes to Brain-derived neurotrophic factor induced protein translation

Transcripts encoding membrane and secreted proteins as well as ER resident proteins have been reported to be present in axons, and the corresponding proteins are synthesized and delivered into the axoplasmic membrane (Willis, Li et al. 2005, Tsai, Bi et al. 2006, Willis, van Niekerk et al. 2007). Initial ultrastructural approaches have identified RER in dendrites (Farah, Liazoghli et al. 2005) but only SER in axons (Tsukita and Ishikawa 1976). However, fluorescence microscopy approaches have identified ER-associated proteins implicated in protein translocation, folding, and post-translational modifications as well as proteins of the Golgi apparatus in distal axons suggesting that axons might contain RER (Merianda, Lin et al. 2009). Our data show that transmembrane proteins such as TrkB as well as N-type Ca^2+^ channels undergo intra-axonal translation in response to BDNF stimulation. Accordingly, we considered whether the axonal growth cones harbors RER, thereby participating in processing of these locally produced membrane and secretory proteins. To address this hypothesis, we transduced motoneurons with lentiviruses encoding the ER marker mCherry-KDEL and co-stained against RPL24 and RPS6 as markers of entirely assembled 80S ribosomes (Fig. 8A). In contrast to unstimulated neurons, a significant number of RPL24/RPS6 co-clusters colocalized with ER in axonal growth cones of stimulated neurons, and this colocalization increased after 10 sec as well as 1 min but dropped again after 30 min BDNF pulse (Fig. 8B). Next, we investigated the role of actin in dynamic assembly of the RER in axon terminals. Interestingly, in line with our previous data shown in Fig. 5L,M and Fig. 6D, CytoD treatment diminished the translocation of ribosomes toward the ER confirming the role of actin in the rapid assembly of RER (Fig. 8C). At last, we examined the attachment of actively translating ribosomes to ER in axonal growth cones by staining mCherry-KDEL expressing neurons against Y10B and eEF2 as depicted in Fig. 8D. Similarly, 10 sec and 1 min stimulations caused an increase in colocalization of Y10B/eEF2 co-clusters with the ER indicating that ribosomes in the elongation stage of translation attach to the ER in axonal growth cones (Fig. 8E). These findings imply that RER exists in the growth cone of developing neurons and thus support a role for ER in processing and surface delivery of axonally synthesized membrane and secretory proteins.

**Figure 8.**
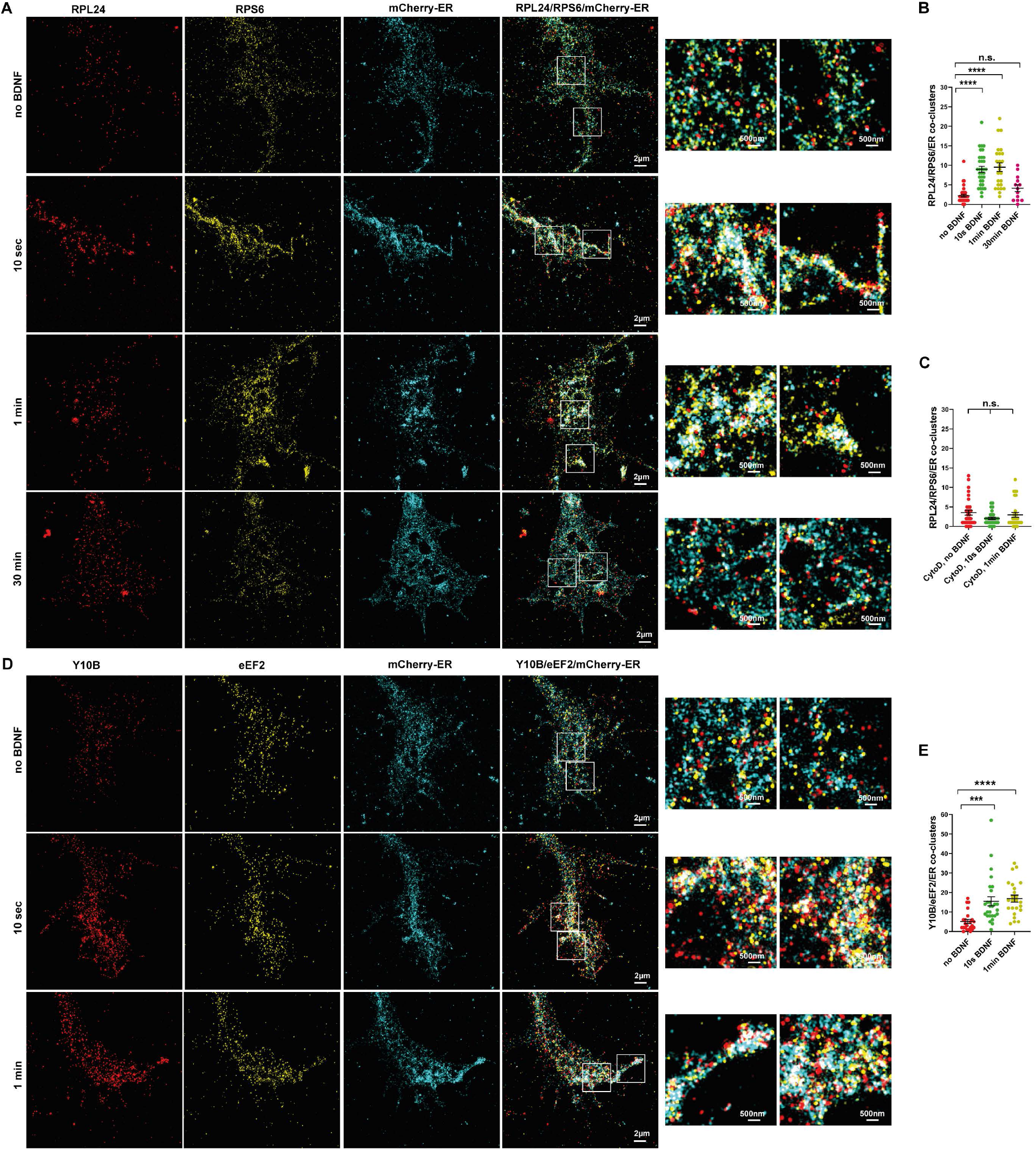
Ribosomes attach to the ER in axonal growth cones and promote local translation in response to BDNF stimulation. (A) BDNF-stimulated motoneurons expressing mCherry-ER were stained against RPL24 and RPS6, and growth cones were imaged by SIM. White squares indicate magnification of ROIs in growth cones. (B) Graph shows co-cluster formation of RPL24/RPS6 with ER within 10 sec and 1 min of stimulation (****, P<0.0001; n=25-33 cells from 3 independent experiments) that dissociate from ER after long stimulation of 30 min (n.s., P=0.598; n=13 from 2 independent experiments). (C) CytoD-treated neurons do not show increased number of RPL24/RPS6/ER co-clusters in growth cones in response to BDNF pulse (n.s., P=0.728; n=30-32 cells from 3 independent experiments). (D) BDNF-stimulated motoneurons expressing mCherry-ER were stained against Y10B and eEF2 and growth cones were imaged by SIM. White squares indicate magnification of ROIs in growth cones. (E) Graph shows co-cluster formation of Y10B/eEF2 with ER within 10 sec and 1 min stimulation (***, P=0.0003; ****, P<0.0001; n=23-27 cells from 3 independent experiments). Graphs are shown in scatter dot plot with mean±SEM. Statistical analyses: one-way ANOVA with Dunn’s post-test.

## Discussion

In neurons, ER forms a highly dynamic network in dendrites and axons. The regulation of axonal ER dynamic movements as well as its function in development and maintenance of synapses have gained emerging interest (Summerville, Faust et al. 2016, De Gregorio, Delgado et al. 2017, Lindhout, Cao et al. 2019). In this study, we developed a technique to visualize and quantify the dynamic movements of ER in axon terminals of cultured motoneurons and show that motoneurons harbor a rough ER in the axonal growth cone, which extends into filopodia and its integrity and dynamic remodeling are regulated mainly by actin and myosin VI.

Contrary to microtubules (Guo, Li et al. 2018), the interaction of ER with actin cytoskeleton especially in the axonal growth cones where presynaptic structures form under specific culture conditions (Jablonka, Beck et al. 2007) has not been investigated yet in much detail. In somatodendritic compartments, kinesin and dynein-based ER sliding along microtubules as well as interaction between STIM1 on the ER and EB1 on the microtubule plus end have shown to mediate ER movements (Waterman-Storer and Salmon 1998, Friedman, Webster et al. 2010). However, studies with cultured cells have revealed that first, nocodazole treatment does not retract ER from the periphery immediately (Terasaki, Chen et al. 1986), second, ER tubules can form in the absence of microtubules (Dreier and Rapoport 2000, Voeltz, Prinz et al. 2006, Wang, Romano et al. 2013) and third, disruption of actin filaments in hippocampal neurons affects Ca^2+^ release from ER in soma (Wang, Mattson et al. 2002). These observations indicate that ER dynamic movements might not depend only on microtubules but also on the actin cytoskeleton. Indeed, the first evidence for the role of actin in ER translocation into dendrites comes from a study by Wagner et al., showing that myosin Va, which is an actin-based motor, mediates ER movement in dendritic spines of Purkinje neurons (Wagner, Brenowitz et al. 2011). Axonal growth cones consist of highly dynamic membrane protrusions, “filopodia and lamellipodia”, and a more stable central region called “growth cone core” (Dent and Gertler 2003). All these domains are transient and undergo constant growth and retraction in shape and structure (Dent, Gupton et al. 2011). In contrast to microtubule-rich core, actin is predominant in filopodia and lamellipodia. Therefore, dynamics of filopodia exceed those in the core (Mallavarapu and Mitchison 1999, Bornschlögl, Romero et al. 2013). In line with that, our time-lapse recordings show that the dynamics of ER differs in growth cone center and periphery. In the core of growth cones, the velocity of ER dynamic movements is much lower than that in actin-rich filopodia. Strikingly, actin depolymerization destroys ER dynamic movements in filopodia of 80% of imaged cells. In addition, two-color live cell imaging data revealed that ER moves along actin in filopodia and F-actin withdrawal results in nearly 99% retraction of ER from filopodia. Finally, pharmacological inhibition of myosin V and particularly of myosin VI significantly reduces the ER movements in filopodia indicating that ER movements in growth cone require myosin VI. These data provide the first direct evidence of an actin/myosin VI dependent ER remodeling and integrity in axon terminals of neurons. ER entry into filopodia suggests a membrane contact site between the ER and the plasma membrane (Wu, Whiteus et al. 2017, Cohen, Valm et al. 2018). This physical tethering could be essential for the surface delivery of newly synthesized receptors, membrane proteins such as voltage gated Ca^2+^ channels, as shown here, or lipids. Importantly, we found that ER movements in the core are regulated by both actin and microtubule cytoskeleton. Previous studies have demonstrated that F-actin and microtubules interplay during neuronal polarization and development (Zhao, Meka et al. 2017, Dogterom and Koenderink 2019, Meka, Scharrenberg et al. 2019). This crosstalk is mediated by macromolecules such as drebrin E, which physically links F-actin and microtubule plus end binding protein or other regulatory proteins such as Rho GTPases, MAP2 and tau (Zhao, Meka et al. 2017, Dogterom and Koenderink 2019). We showed that, knockdown of drebrin A significantly decreases the dynamic remodeling of axonal ER indicating a role for this protein in the coordinated actin/microtubule regulation of ER dynamics in axonal growth cones.

Subcellular control of axon growth, synapse differentiation and plasticity depend on regulatory mechanisms that ensure processing of local information and subsequent responses on a time scale of seconds. mRNA localization and local protein synthesis are conserved mechanisms that modify axonal proteome at a fast temporal and spatial scale, thereby maintaining plasticity capacity of axonal synapses (Holt, Martin et al. 2019, Fernandopulle, Lippincott-Schwartz et al. 2021). In addition to cytoplasmic proteins, mRNAs encoding secreted and membrane proteins have been identified in axonal transcriptomes (Willis, van Niekerk et al. 2007, Saal, Briese et al. 2014, Poulopoulos, Murphy et al. 2019). On-site synthesis and glycosylation of such membrane proteins would require additional RER and Golgi machinery in the distal axon. However, previous studies have suggested that in axons ER is considered to be devoid of ribosomes and thus no direct evidence of its potential functions related to local protein synthesis has been reported yet (Tsukita and Ishikawa 1976, Krijnse-Locker, Parton et al. 1995, Horton and Ehlers 2003). Interestingly, Merianda et al., 2009 reported glycosylation patterns of transmembrane proteins being locally translated in axons of sensory neurons (Merianda, Lin et al. 2009). We tested the effects of a BDNF pulse in cultured motoneurons that were then investigated with super-resolution fluorescence imaging by SIM. This approach revealed that stimulation of motoneurons triggers the assembly of 80S ribosomes and initiates the translation in axon terminals within 10 sec. Translationally active ribosomes attach to the ER in axonal growth cones and local production of new proteins including transmembrane proteins such as TrkB and α-1β subunits of N-type Ca^2+^ channels becomes detectable within 1 min poststimulation. Contrary to axons, the response of soma to BDNF/TrkB activation was considerably slower and happened in longer time scales of several minutes. These distinctive response rates could be due to different intracellular levels of second messengers such as cAMP (Ming, Song et al. 1997). It remains unknown whether similar or distinct signaling pathways downstream to BDNF/TrkB, such as mTOR, MAPK and PI3K pathways, are involved in activation of ribosomes and translation initiation in the growth cone and its counterparts. The rate of dynamic local translation at axonal terminals of cultured motoneurons appears relatively high. The fast and dynamic translocation of ribosomal subunits to ER that rapidly forms a rough ER in axon terminals, thus provides the basis for post-translational processing of locally synthesized proteins in response to extracellular stimuli. Intriguingly, 30 min after stimulation, 80S ribosomal subunits disassemble again and further dissociate from the ER indicating that ribosomes only transiently associate with axonal ER in response to stimuli and explains why previous studies have failed to detect RER in axons using unstimulated neurons. Moreover, our data show that the rapid assembly of 80S ribosomes as well as their translocation toward the RER are disrupted upon pharmacological inhibition of actin assembly suggesting involvement of an actin regulatory mechanism.

In summary, this work identifies a novel function for axonal ER in regulation of stimulus-induced local translation and discloses a mechanism for dynamic regulation of ER in axonal growth cones by a drebrin A-mediated actin and microtubule crosstalk.

## Materials and methods

### Animals

TrkB knockout mice (Rohrer, Korenbrot et al. 1999) were obtained from the University of California, Davis (MMRRC: 000188, B6; 129S4-Ntrk2 < tm1Rohr >) and maintained on a C57Bl/6 background. CD1 mice were used for motoneuron cell cultures from WT mice. All mice were housed in the animal facility of the Institute of Clinical Neurobiology, University Hospital of Wuerzburg. All mouse procedures were performed according to the regulations on animal protection of the German federal law and of the Association for Assessment and Accreditation of Laboratory Animal care, approved by the local authorities.

### Culture of embryonic mouse motoneurons

Embryonic mouse motoneuron culture was performed as previously described (Wiese, Herrmann et al. 2009). Lumbar spinal cords were dissected from E13.5 CD1 mouse embryos, digested with trypsin (Worthington), triturated and then transferred onto a panning plate coated with anti-p75 antibody (MLR2, Abcam) for enrichment of motoneurons. For lentiviral transduction, pSIH-mCherry-KDEL construct expressing lentiviruses were added to the suspension of motoneurons before plating on polyornithine-(PORN) and human merosin-coated (CC085, Merck-Millipore) plates. Motoneurons were cultured in NB medium supplemented with 500 µM Glutamax, 2% heat-inactivated horse serum (Gibco), 2% B27 (Thermo Fisher Scientific) and 5 ng/ml BDNF for 5 to 6 days in a humidified CO2 incubator at 37°C. Medium was changed 24 h after plating and then every other day. Merosin consists of Laminin211 (α2β1γ1), enriched in extrasynaptic basal lamina and Laminin221 (α2β2γ1), which is specifically expressed at the cleft of NMJs and regulates formation, maturation and maintenance of NMJs (Fox, Sanes et al. 2007, Rogers and Nishimune 2017). Thus, culturing motoneurons on merosin induces the maturation of presynaptic structures in axon terminals. Compartmentalized motoneuron cultures were prepared as described previously (Saal, Briese et al. 2014). To drive axonal growth into the axonal compartment, 20 ng/ml BDNF and CNTF were added into the axonal compartments, while only 5 ng/ml CNTF was added into the somatodendritic compartments.

### Cloning and lentivirus production

For cloning of pSIH-mCherry-KDEL, mCherry was first amplified by PCR using a commercially available plasmid as template (primers: forward: 5’-ACTCGTCGACGTGAGCAAGGGCGAGGAGGAT-3’; reverse: 5’-GAATGCGGCCGCCTTGTACAGCTCGTCCATGCC-3’), and cut by SalI and NotI enzymes, followed by insertion into the pCMV-ECS2-CMV-myc-ER vector backbone containing a KDEL sequence. The assembled mCherry-KDEL fragment was amplified by PCR from the above vector (primers: forward: 5’-TTTGACCTCCATAGAAGATTCCACCATGGGATGGAGCTG-3, reverse: 5’-TGTAATCCAGAGGTTGATTGCTACAGCTCGTCCTTCTCG-3’), and the product was inserted into a lentiviral expression vector (pSIH-H1) containing CMV promotor using NEBbuilder HIFI kit (NEW ENGLAND Biolabs). To co-express GFP-actin and mCherry-KDEL, a previously described GFP-actin construct was used (Sivadasan, Hornburg et al. 2016). Both mCherry-KDEL and GFP-actin were first amplified by PCR and purified fragments were inserted into a pSIH-CMV-IRES vector. To achieve knockdown of drebrin isoforms, shRNAs targeting mouse drebrin A and shRNAs targeting both drebrin A and E were cloned into a modified version of pSIH-H1 shRNA vector (System Biosciences) containing eGFP according to the manufacturer’s instructions. The following antisense oligonucleotides were used: shDrebrin A: 5’-GTCCGTACTGCCCTTTCATAA-3’, shDrebrin A+E: 5’-GGCTGTGCTAACCTTCTTAAT-3’. Empty pSIH-H1 expressing eGFP was used as shControl. Lentiviruses were produced in HEK^293T^ cells using pCMV-VSVG and pCMVΔR8.91 helper plasmids (Subramanian, Wetzel et al. 2012). HEK^293T^ cells were transfected with calcium-phosphate reagents, and viral supernatants were harvested after 47 h by ultracentrifugation. NSC34 cells were used for virus titer test.

### Quantitative qRT-PCR

Total RNA was extracted separately from somatodendritc and axonal compartments and reverse-transcribed using random hexamers and Superscript III Reverse transcriptase enzyme (18080044; Invitrogen), as described previously (Moradi, Sivadasan et al. 2017). cDNA was purified using QIAGEN II purification kit (20021). For qRT-PCR, Luminaris HiGreen qPCR Master Mix (Thermo Fisher Scientific) was applied on a lightCycler® 96 thermal cycler (Roche). Histone H1f0 transcripts were absent in RNA fractions obtained from axonal compartments, confirming the purity of mRNA preparation from these compartments. GAPDH was used for data normalization. The following primers were used for qRT-PCR: TrkB: 5’-CGGGAGCATCTCTCGGTCTAT-3’ (forward) and 5’-CTGGCAGAGTCATCGTCGTTG-3’ (reverse); Gapdh: 5’-AACTCCCACTCTTCCACCTTC-3’ (forward), and 5’-GGTCCAGGGTTTCTTACTCCTT-3’ (reverse); and histone H1f0, 5’-CCCAAGTATTCAGACATGAT-3′ (forward), and 5’-CGCTTGATGGACAACT-3’ (reverse). To assess knockdown efficiency of shDrebrin A and shDrebrin A+E lentiviruses, primary motoneurons were transduced with knockdown lentiviral constructs and cultured for 7 days. Cells were first washed 2 x with RNAase free PBS and then lysed in RNA lysis buffer. RNA was extracted using NucleoSpin RNA extraction kit (Macherey-Nagel) and reverse-transcribed with random hexamers using First Strand cDNA Synthesis Kit (Thermo Fisher Scientific). Reverse transcription reactions were diluted 1:5 in water and proceeded with qRT-PCR. Following primers were used for qRT-PCR reactions: for drebrin A: forward: 5’-CCTGATAACCCACGGGAGTT-3’, reverse: 5’-GGAAGAGAGGTTTGGGGTGC-3’; for drebrin E: forward: 5’-CCCACGGGAGTTCTTCAGACA-3’, reverse: 5’-TCCAGGTGGCTGCATGGGAGGGAG-3’).

### BDNF stimulation and immunocytochemistry of cultured motoneurons

At DIV6, motoneurons were fixed with 4% PFA for 10 min at RT and permeabilized with 0.1% or 0.3% Triton X-100 for 10 min, followed by 3 x wash with PBS. After incubation with block solution (10% donkey serum and 2% BSA in PBS) at RT for 1 h, primary antibodies were added and incubated at 4°C overnight. On the second day, motoneurons were washed trice with PBS and incubated with secondary antibodies for 1 h, followed by another 3x wash. Aqua Poly/Mount (18606-20, Polysciences) was used for embedding. For β-actin staining, motoneurons were permeabilized with ice-cold methanol for 5 min at -20°C. For Cav2.2 staining, motoneurons were fixed with 4% PFA for 5 min at RT and permeabilized with 0.1% Triton X-100 in PBS for 5 min at RT. For the BDNF stimulation experiments, motoneurons were first cultured with 5 ng/ml BDNF. At DIV5, medium was completely aspired and cells were washed twice with NB medium to completely remove BDNF from all the surfaces. Cells were maintained in NB medium supplemented with 2% HS and 2% B27 overnight in the absence of neurotrophic factors. On the next day, BDNF stimulation was conducted by directly adding 40 ng/ml BDNF into the cell culture medium and cultures were kept on a 37°C hotplate. For different stimulation times, BDNF containing medium was removed after 10 sec, 1 min and 10 min stimulation and 4% PFA was directly added onto cells. For the no BDNF control group, the same amount of NB medium without BDNF was added. The following primary antibodies were used: monoclonal mouse anti-α-Tubulin (T5168, Sigma-Aldrich; 1:1000), polyclonal goat anti-TrkB (AF1494, Bio-Techne Sales Corp; 1:500), monoclonal mouse anti-rRNA (Y10b) (MA116628, Thermo Fisher Scientific; 1:500), polyclonal goat anti-ribosomal protein L8 (SAB2500882, Sigma-Aldrich; 1:500), monoclonal mouse anti-β-Actin (GTX26276, GeneTex; 1:1000), polyclonal rabbit anti-Tau (T6402,Sigma-Aldrich;1:1000), polyclonal rabbit anti-eEF2 (23325, Cell Signaling Technology; 1:50), monoclonal mouse anti-S6 ribosomal protein (MA515123, Thermo Fisher Scientific; 1:100), polyclonal rabbit anti-RPL24 (PA530157, Thermo Fisher Scientific; 1:500), polyclonal guinea pig antiserum RFP (390004, Synaptic Systems; 1:500), polyclonal guinea pig Ca^2+^ channel N-type alpha-1B channel (152305, Synaptic Systems, 1:250) and monoclonal rat anti-mCherry (M11217, Thermo Fisher Scientific; 1:1500). Secondary antibodies: donkey anti-mouse IgG (H+L) (Alexa Fluor 488; A21202,Life Technologies;1:500), donkey anti-rabbit IgG (H+l) AffiniPure (Alexa Fluor 488; 711-545-152, Jackson ImmunoResearch; 1:500), donkey anti-guinea pig IgG (H+L) AffiniPure (Cy3; 706-165-148: Jackson ImmunoResearch; 1:500), donkey anti-rat IgG (H+L) AffiniPure (Cy3; 712-165-150; Jackson ImmunoResearch), donkey anti-rabbit IgG (H+L) AffiniPure (Cy3; 711-165-152; Jackson ImmunoResearch), donkey anti-goat IgG (H+L) AffiniPure (Cy3;705-165-147: Jackson ImmunoResearch), donkey anti-mouse IgG (H+L) (Cy3; 715-165-151; Jackson ImmunoResearch; 1:500), donkey anti-goat IgG (H+L) AffiniPure (Alexa Fluor 647; 705-605-003; Jackson ImmunoResearch), donkey anti-mouse IgG (H+L) highly cross-adsorbed (Alexa Fluor 647; A31571; Invitrogen), donkey anti-rabbit IgG (H+L) AffiniPure (Cy5; 711-175-152; Jackson ImmunoResearch). F-actin was labeled with Alexa Fluor647-conjugated Phalloidin (A22287; Invitrogen). All secondary antibodies were diluted 1:500 in TBST.

### Puromycin experiments

Motoneurons were incubated with 10 µg/ml puromycin for 10 min and BDNF stimulation was carried out as described in the above section. Cells were then fixed and immunostained using anti-puromycin immunostaining. Nocodazole was used to disrupt microtubule-dependent axonal transport and anisomycin was used as translational inhibitor. Cells were treated with 10 µM nocodazole for 2 h and 100 ng/ml anisomycin for 1 h prior to as well as during puromycin incubation followed by BDNF stimulation. Primary and secondary antibodies used were: monoclonal mouse anti-puromycin (clone 12D10, MABE343, Merck Millipore; 1:1000), polyclonal rabbit anti-Tau (T6402, Sigma-Aldrich; 1:1000), donkey anti-mouse IgG (H+L) (Alexa Fluor 488; A21202, Life Technologies; 1:500), and donkey anti-rabbit IgG (H+L) AffiniPure (Cy3; 711-165-152; Jackson ImmunoResearch).

### Image acquisition and data analysis

Image acquisition was done with a standard Olympus Fluoview 1000 confocal system with a 60x NA 1.35 oil objective. Structured illumination microscopy (SIM) imaging was performed on a commercial ELYRA S.1 microscope (Zeiss AG). The setup is equipped with a Plan-Apochromat 63x NA 1.40 immersion-oil based objective and four excitation lasers, a 405 nm diode (50 mW), a 488 nm OPSL (100 mW), a 561 nm OPSL (100 mW) and a 642 nm diode laser (150 mW). 3D reconstruction was performed using Imaris software. For quantification of immunofluorescence signals, mean gray values of images were measured from unprocessed raw data after background subtraction using ImageJ-win64. For quantification of SIM data represented in Fig.4 and 6, co-clusters of RPL24/RPS6, Y10B/eEF2 as well as RPL24/RPS6/ER and Y10B/eEF2/ER were counted manually. For this, maximum projections of single SIM channels (RPL24, RPS6 and mCherry-ER or Y10B, eEF2 and mCherry-ER channels) were first created using ImageJ. An automatic linear adjustment of contrast and brightness was applied to the whole z-projection image of each single channel and these were merged into a RGB image. Co-clusters were defined as overlapping dots from RPL24, RPS6 and mCherry-ER channels or Y10B, eEF2 and mCherry-ER channels on the RGB image which had a diameter of more than 350 nm. In addition, colocalization of RPL24/RPS6 as well as Y10B/eEF2 shown in Fig. 4 was assessed by Pearson’s correlation coefficient using Coloc2 plugin of Fiji. GraphPad Prism 9 software was used for all statistical analyses. Data are shown in scatter dot plots with mean ± SEM, unless otherwise mentioned. For a better visibility, linear contrast enhancement was applied to all representative images using Adobe Photoshop.

### Live cell Imaging and data quantification

Approximately, 40,000 motoneurons were transduced with lentivirus expressing pSIH-mCherry-KDEL or pSIH-GFP-actin-IRES-mCherry-KDEL and cultured on PORN/ merosin-coated 35 mm high µ-dishes (81156, IBIDI) for 6 days. For pharmacological treatments, cytochalasin D (Sigma-Aldrich; C2618), nocodazole (Sigma-Aldrich; M1404), (-)-blebbistatin (Sigma-Aldrich; B0560), MyoVin-1 (Calbiochem; 475984) and 2,4,6-triiodophenol (Sigma-Aldrich; 137723) were dissolved and diluted in DMSO (PanReac AppliChem). Cells were treated at the following concentration and incubation time before live cell imaging: 1 µg/ml cytochalasin D for 30 min; 10 µM nocodazole for 2 h; 5 µM (-)-blebbistatin for 15 min; 30 µM MyoVin-1 for 15 min and 1 µM 2,4,6-triiodophenol for 15 min. For live cell imaging, cells were washed 2x with pre-warmed Tyrode’s Solution (125 mM NaCl, 2 mM KCl, 2 mM CaCl2, 2 mM MgCl2, 30 mM glucose, 25 mM HEPES, pH 7.4) and subsequently imaged in 2 ml Tyrode’s Solution. Imaging was performed using an inverted epifluorescence microscopy (TE2000; Nikon) that was equipped with a perfect focus system, heated stage chamber (TOKAI HIT CO, LTD) at 37 °C, 5% CO2 and 60x 1.4 NA objective. Time series were captured at a speed of 2 sec per frame over 15 min. 12-bit images of 1.024 × 1.024 pixels were acquired with an Orca Flash 4.0 V2 camera (Hamamatsu Photonics), controlled by Nikon Element image software. For quantification of ER dynamics, Image Correlation Spectroscopy (ICS) (Wiseman 2015) was implemented in python. This approach was used as described previously by Wiseman et,. al. Briefly, molecular movements of ER or molecular transport of other organelles are determined based on flow or diffusion in an image time series 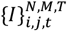, with N, M, T indicating the two dimensions in space and the time dimension respectively. *i, j, t* denote the corresponding running indices the basic workflow starts by defining a space-time window, i.e., a subspace of *K = L =* 10 *p*_*x*_ over Δ*t = 10* consecutive frames. This subspace is rasterized over the image with a sampling rate of Δi = Δj = 4. For each of those samples a correlation window 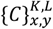 is computed, describing the overlap of signal for a given shift *x, y* per time frame. The maximum of this spectrum is the point of maximum correlation and indicates a shift of signal from one image to this next, by (Δi,Δj) *=* (*x*_*max*_ −*K*/2, *y*_*max*_−*L*/2) _(1)_. To achieve sub-pixel accuracy a 2D Gaussian function is fitted to the correlation spectrum using the Levenberg-Marquardt-Algorithm. The initial parameters were chosen as amplitude *A = max* (*C*), center coordinates in *x, x*_0_ *= K*/2 _(2)_, center coordinates in *y, y*_0_ *= L*/2 _(3)_, standard deviation in *x, σ*_*x*_ *= K*/4 _(4)_, standard deviation in *y, σ*_*y*_ *= L*/4 _(5)_. An additional parameter *θ* rotating the coordinate system by *θ* was initialized as 0. The fit results for *x*_0_ and *y*_0_ indicate a directed shift of signal. A larger *σ*_*x*_ or *σ*_*y*_ indicates an overall lower correlation, i.e., the signal losing its shape. To compute overall dynamics, three distinct subspaces are classified: (i) subspaces with a mean intensity less than 10% of the maximum intensity are considered noise. (ii) Subspaces with a mean intensity that is larger than 40% of the maximum intensity are classified as core. (iii) Subspaces with an intensity in between is considered as filopodia. The overall dynamics value for a class is the mean dynamics value of all subspaces. Kymograph-analysis of ER dynamics was carried out using Image J as described previously (Moradi, Sivadasan et al. 2017). Briefly, a Z-projection image was made from all the frames of a live cell image, followed by the creation of a multiple kymograph along a line drawn tracking the ER movements in filopodia or core. Distance changed per movement and frequency of movements over 15 min were calculated from multiple kymographs and plotted in a graph. Growth cones without detectable ER movements were classified as failure filopodia or core ER movement.

### Western blotting

For Western blot analysis, approximately 300,000 motoneurons were plated onto a PORN/merosin-coated 24-well cell culture dish. At DIV6, BDNF-deprived neurons were stimulated with 40 ng/ml BDNF for 10 sec, 30 sec, 1 min and 10 min and cells were lysed directly in 2x Laemmli buffer (125 mM Tris, pH 6.8, 10% SDS, 50% glycerol, 25% β-mercaptoethanol and 0.2% bromophenol blue). The protein lysates were boiled at 99°C for 10 min. Protein extracts were separated using 4-12% Gradient SDS-PAGE gels. Proteins were then blotted onto nitrocellulose membranes and primary antibodies were incubated on a shaker at 4°C overnight. Primary antibodies were washed with TBST and secondary antibodies were incubated at RT for 1 h, washed in TBST and developed using ECL systems (GE Healthcare). The primary antibodies used are polyclonal goat anti-TrkB (AF1494, Bio-Techne Sales Corp; 1:500) and monoclonal mouse anti-α-tubulin (T5168, Sigma-Aldrich; 1:5000). The secondary antibodies used are peroxidase AffiniPure donkey anti-rabbit IgG (H+L) (711-035-152, Jackson ImmunoResearch; 1:10,000), peroxidase AffiniPure donkey anti-goat IgG (H+L) (705-035-003, Jackson ImmunoResearch; 1:10,000) and peroxidase AffiniPure goat anti-mouse IgG (H+L) (115-035-146, Jackson ImmunoResearch; 1:10,000).

## Supporting information

Dynamic movements of ER and plasma membrane in the axonal growth cone of motoneurons

Co-movements of ER and actin in axonal growth cone filopodia

ER growth and retraction in axonal growth cone filopodia do not completely depend on actin movements.

Dynamic movements of ER in the core of the axonal growth cone of untreated motoneurons

Dynamic movements of ER in filopodia of the axonal growth cone of untreated motoneurons

Dynamic movements of ER in the core of the axonal growth cone of CytoD-treated motoneurons

Dynamic movements of ER in the core of the axonal growth cone of nocodazole-treated motoneurons

Dynamic movements of ER in filopodia of the axonal growth cone of CytoD-treated motoneurons

Dynamic movements of ER in filopodia of the axonal growth cone of nocodazole-treated motoneurons

Dynamic movements of ER in axonal growth cones of neurons treated with myosin II inhibitor

Dynamic movements of ER in axonal growth cones of neurons treated with myosin V inhibitor

Dynamic movements of ER in axonal growth cones of neurons treated with myosin VI inhibitor

Dynamic movements of ER in axonal growth cones of neurons transduced with shDrebrin A

Dynamic movements of ER in axonal growth cones of neurons transduced with shDrebrin A+E

## Abbreviation

BDNF: Brain-derived neurotrophic factor
TrkB: Tropomyosin receptor kinase B

## Abbreviation table

ALS: Amyotrophic Lateral Sclerosi
BDNF: Brain-derived Neurotrophic Factor
ER: Endoplasmic Reticulum
HSP: Hereditary Spastic Paraplegia
NMJs: Neuromuscular Junctions
SIM: Structured Illumination Microscopy
TVA: Transverse Abdominal muscle
TrkB: Tropomyosin receptor kinase B

## Acknowledgments

We thank Markus Behringer for assistance with SIM, Regine Sendtner for animal breeding, Hildegard Troll for lentiviral production and Dr. Robert Blum for helpful discussions.

## Funding

Chunchu Deng was supported by a grant provided from PicoQuant. PicoQuant was not involved in design, performing or analysis of data. This paper received further support from the Deutsche Forschungsgemeinschaft (DFG), Grant Se697/7-1 for MS, and Deutsche Forschungsgemeinschaft (DFG), Grant JA1823/3-1 for SJ.

The authors declare no competing financial interests.

## Authors’ contributions

CCD, MM and M. Sendtner conceived the project, designed the experiments and wrote the manuscript. CCD performed all the experiments including live cell imaging of ER dynamics, motoneuron cultures, ICC, IHC, SIM and confocal microscopy and data analysis. MM performed qRT-PCR and compartmentalized motoneuron cultures. SR and SD performed analysis of ER dynamic movements. MM, LH, PL and SJ helped with confocal image acquisition and data analysis. CHJ constructed drebrin knockdown plasmids. PL provided 3D SIM images. M. Sauer gave support with super resolution microscopy. LH provided additional data for discussions.

## Notes

### Competing Interest Statement

The authors have declared no competing interest.

