## Supplementary Figures for "Dynamic remodeling of ribosomes and endoplasmic reticulum in axon terminals of motoneurons"

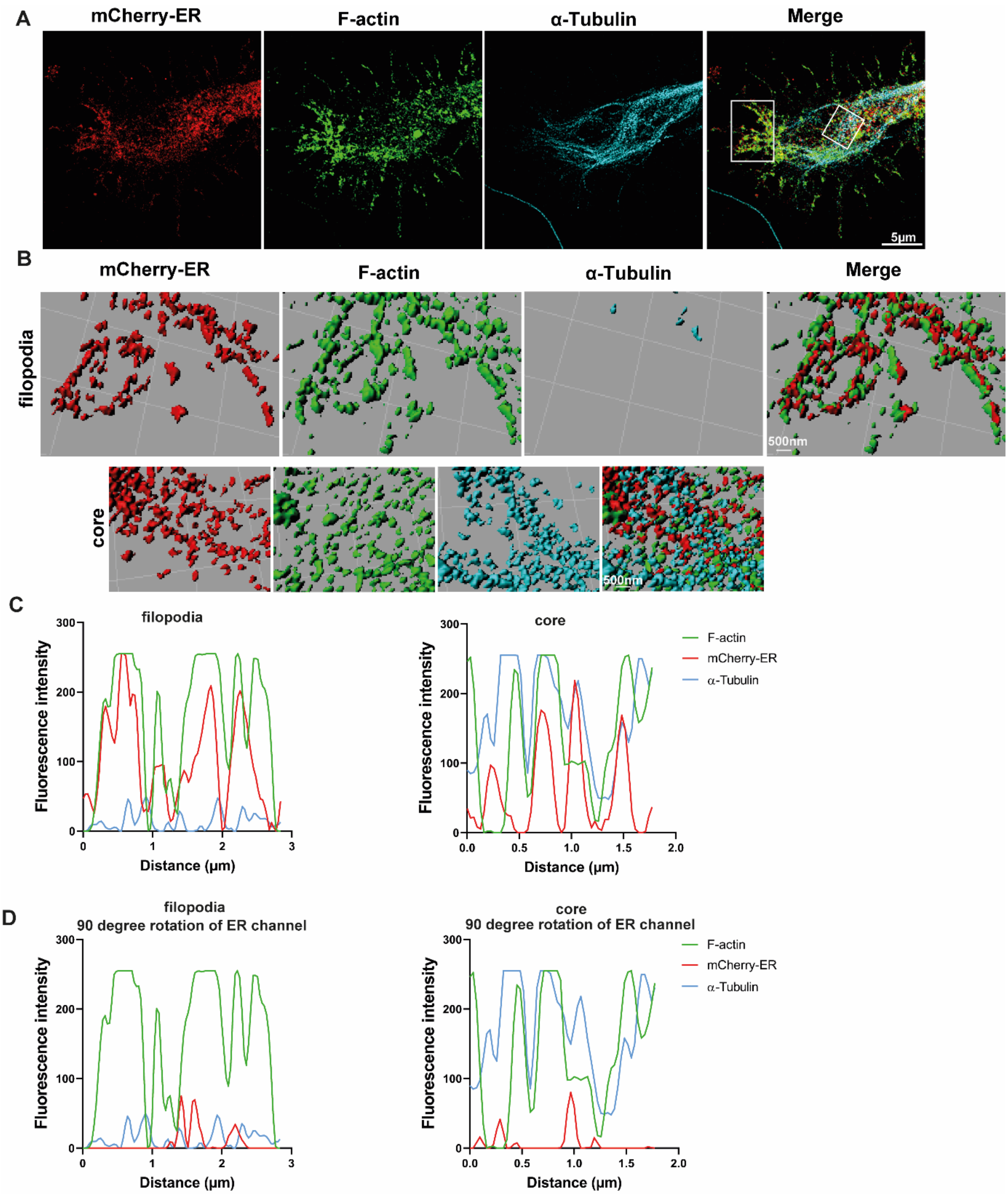


Figure S1.

**ER is present in the growth cone of cultured motoneurons and enters filopodia.** (A) Motoneurons were transduced with a lentiviral construct expressing mCherry-KDEL (mCherry-ER) to visualize ER in the growth cone. To determine growth cone boundaries, motoneurons were immunostained against F-actin and α-Tubulin in addition to mCherry. Images captured by SIM show that mCherry-ER is present in both core and filopodia sub-regions of the growth cone. (B) Representative 3D reconstruction of the overlap image shown in A. White square boxes indicate magnification of ROIs in filopodia and core. (C) Representative line scan diagrams showing overlap of mCherry-ER (red), F-actin (green) and α-Tubulin (blue) channels in growth cone filopodia or core. (D) As control, the ER channel was rotated 90 degree and the colocalization of ER with F-actin or α-Tubulin was evaluated by line scan.

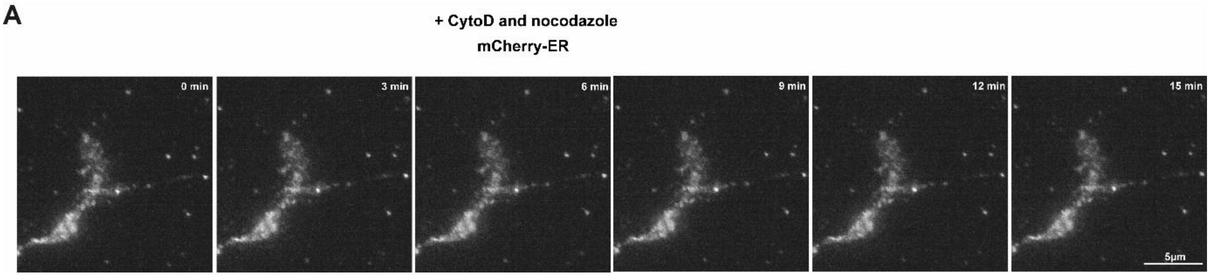


Figure S2.

**Treatment of motoneurons with CytoD and nocodazole disturbs ER dynamic movements in the axonal growth cone.** (A) Representative time-lapse images of motoneurons expressing mCherry-ER that were treated with both nocodazole and CytoD. Almost no more ER movements were detectable.

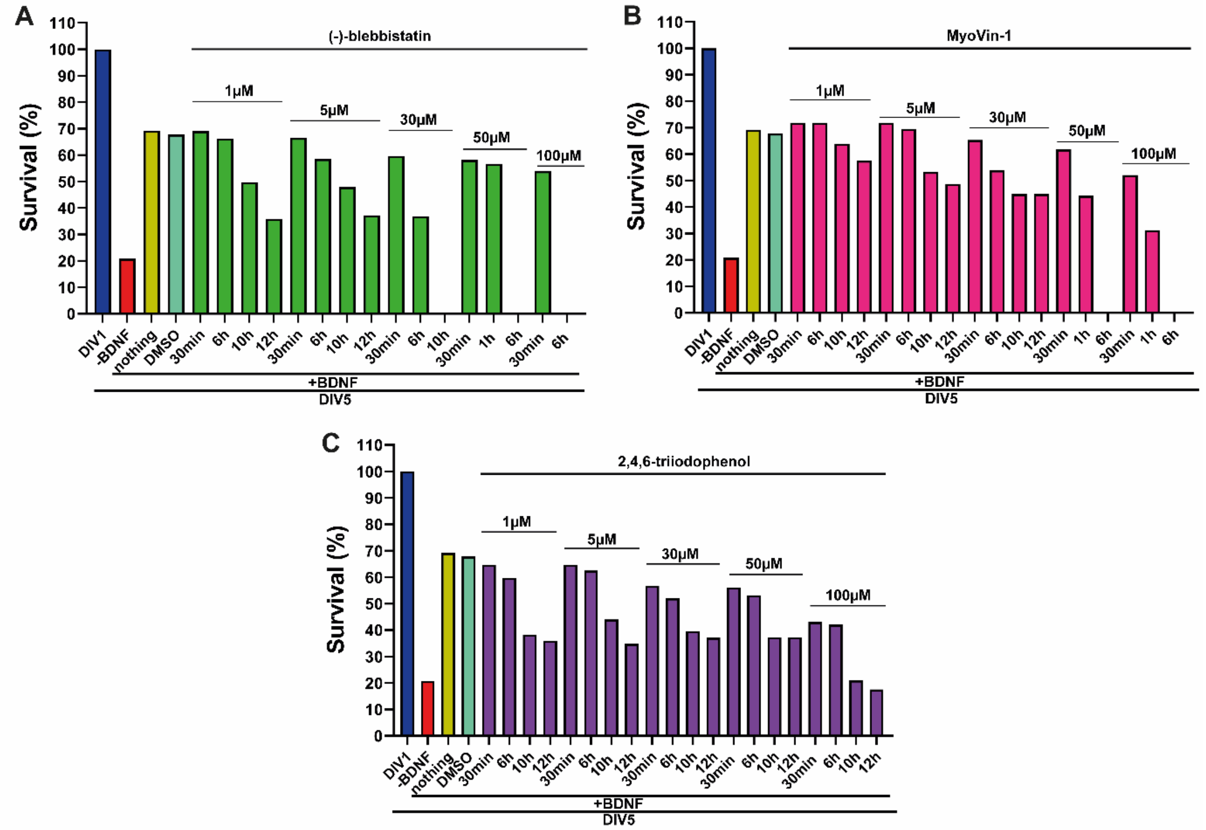


Figure S3.

**Survival assay of motoneurons treated with myosin II, V and VI inhibitors.** Cultured motoneurons were incubated with different doses of myosin inhibitors, ranging from 1-100 µM and incubated for 30 min up to 12 h. (A) Graphs show % of survival of motoneurons incubated with (A) myosin II inhibitor ((-)-blebbistatin), (B) myosin V inhibitor (MyoVin-1), and (C) myosin VI inhibitor (2,4,6-triiodophenol).

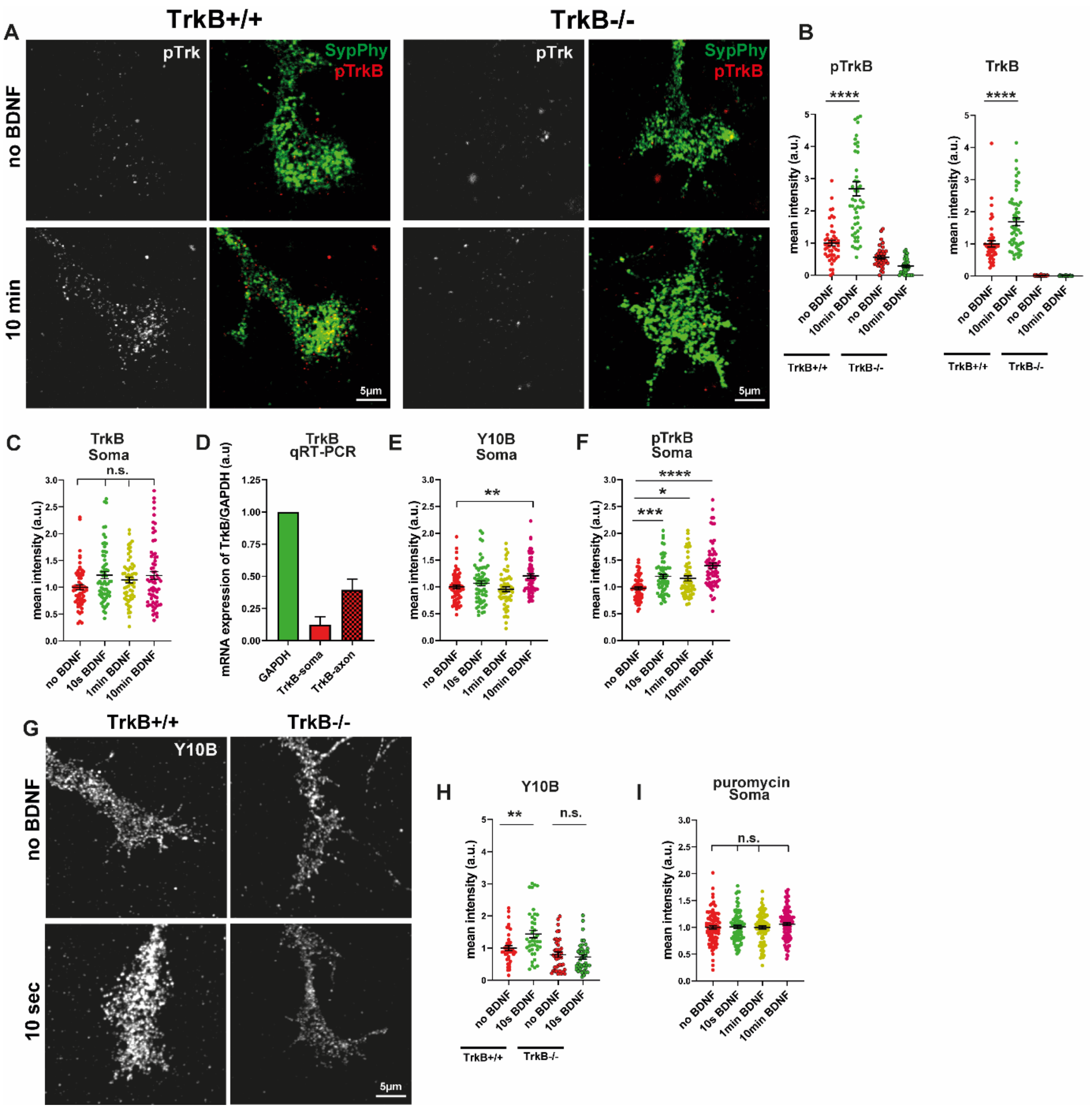


Figure S4.

**BDNF-induced translation initiation is delayed in soma of cultured motoneurons.** (A) Specificity of used antibodies against pTrkB and TrkB was assessed by ICC using motoneurons from wildtype and TrkB knockout mice that received a 10 min BDNF pulse. Representative confocal images of growth cones were stained for pTrkB and synaptophysin. (B) Mean intensities of pTrkB and TrkB were quantified and plotted in a graph (*TrkB+/+*: ****, P<0.0001; n=47-52 cells and *TrkB-/-*: n=34-37 cells from 1 experiment). (C) Graph shows mean intensities of TrkB in soma of BDNF-stimulated motoneurons (n.s., P≥0.1208; n=57-67 cells from 3 independent experiments). (D) Graph shows mRNA levels of TrkB relative to mRNA levels of GAPDH detected by qRT-PCR in somatodendritic as well as axonal compartments of motoneurons cultured in microfluidic chambers. (E) Graph shows mean intensities of Y10B in soma of BDNF-stimulated motoneurons. Increase of mean intensity for Y10B was not observed before 10 min poststimulation (**, P=0.0042; n=57-67 cells from 3 independent experiments). (F) Graph shows mean intensities of pTrkB in soma of BDNF-stimulated motoneurons. Immunoreactivity of pTrkB increases significantly within 10 sec stimulation (***, P=0.0004; n=63-64 cells from 3 independent experiments), as well as 1min and 10min (*, P=0.0378; ****, P<0.0001; n=57-67 cells from 3 independent experiments). (G) Representative confocal images of growth cones of wildtype and TrkB knockout BDNF-stimulated motoneurons stained with Y10B. (H) Mean intensities of Y10B do not increase upon 10 sec BDNF stimulation in TrkB knockout (n.s., P=0.6019; n=40 cells from 2 independent experiments) neurons compared to WT (**, P=0.0025; n=40-41 cells from 2 independent experiments). (I) Graph shows mean intensities of puromycin in soma of BDNF-stimulated motoneurons. Puromycin immunoreactivity is not altered in soma of stimulated neurons (n.s., P≥0.7006; n=83-112 cells from 3 independent experiments). All data are normalized to no BDNF group of the corresponding genotype. Graphs are shown in scatter dot plot with mean±SEM. Statistical analyses: One-way ANOVA with Dunn's post-test in C, E, F and I, and by Mann Whitney test in B and H.

Movie 1.

**Dynamic movements of ER and plasma membrane in the axonal growth cone of motoneurons.** Motoneurons were transduced with lentiviruses expressing mCherry-ER and cell volume marker GFP and imaged using an epifluorescence microscope for 8 min at 2 sec intervals to visualize ER movements in growth cone filopodia. Related to Fig. 1.

Movie 2.

**Co-movements of ER and actin in axonal growth cone filopodia.** Motoneurons expressing mCherry-ER and GFP-actin were imaged using an epifluorescence microscope for 8 min at 2 sec intervals to visualize the interaction of ER and actin in growth cone filopodia. Related to Fig. 1.

Movie 3.

**ER growth and retraction in axonal growth cone filopodia do not completely depend on actin movements.** Motoneurons expressing mCherry-ER and GFP-actin were imaged using an epifluorescence microscope for 8 min at 2 sec intervals. In some filopodia, only ER but not actin collapses. Related to Fig. 1.

Movie 4.

**Dynamic movements of ER in the core of the axonal growth cone of untreated motoneurons.** Motoneurons expressing mCherry-ER were imaged using an epifluorescence microscope for 15 min at 2 sec intervals to visualize ER in the growth cone. Related to Fig. 2.

Movie 5.

**Dynamic movements of ER in filopodia of the axonal growth cone of untreated motoneurons.** Motoneurons expressing mCherry-ER were imaged for 15 min at 2 sec intervals to visualize ER dynamic movements in growth cone filopodia using an epifluorescence microscope. Related to Fig. 2.

Movie 6.

**Dynamic movements of ER in the core of the axonal growth cone of CytoD-treated motoneurons.** Motoneurons expressing mCherry-ER were treated with CytoD and ER dynamic movements were imaged for 15 min at 2 sec intervals using an epifluorescence microscope. Related to Fig. 2.

Movie 7.

**Dynamic movements of ER in the core of the axonal growth cone of nocodazole-treated motoneurons.** Motoneurons expressing mCherry-ER were treated with nocodazole and ER dynamic movements were imaged for 15 min at 2 sec intervals using an epifluorescence microscope. Related to Fig. 2.

Movie 8.

**Dynamic movements of ER in filopodia of the axonal growth cone of CytoD-treated motoneurons.** Motoneurons expressing mCherry-ER were treated with CytoD and ER dynamic movements were imaged for 15 min at 2 sec intervals using an epifluorescence microscope. Related to Fig. 2.

Movie 9.

**Dynamic movements of ER in filopodia of the axonal growth cone of nocodazole-treated motoneurons.** Motoneurons expressing mCherry-ER were treated with nocodazole and ER dynamic movements were imaged for 15 min at 2 sec intervals using an epifluorescence microscope. Related to Fig. 2.

Movie 10.

**Dynamic movements of ER in axonal growth cones of neurons treated with myosin II inhibitor.** Motoneurons expressing mCherry-ER were treated with myosin II inhibitor ((-)-blebbistatin) and ER dynamic movements were imaged for 15 min at 2 sec intervals using an epifluorescence microscope. Related to Fig. 3.

Movie 11.

**Dynamic movements of ER in axonal growth cones of neurons treated with myosin V inhibitor.** Motoneurons expressing mCherry-ER were treated with myosin V inhibitor (MyoVin-1) and ER dynamic movements were imaged for 15 min at 2 sec intervals using an epifluorescence microscope. Related to Fig. 3.

Movie 12.

**Dynamic movements of ER in axonal growth cones of neurons treated with myosin VI inhibitor.** Motoneurons expressing mCherry-ER were treated with myosin VI inhibitor (2,4,6-triiodophenol) and ER dynamic movements were imaged for 15 min at 2 sec intervals using an epifluorescence microscope. Related to Fig. 3.

Movie 13.

**Dynamic movements of ER in axonal growth cones of neurons transduced with shDrebrin A.** Motoneurons were transduced with shDrebrin A and mCherry-ER lentiviruses and ER dynamic movements were imaged for 15 min at 2 sec intervals using an epifluorescence microscope. Related to Fig. 4.

Movie 14.

**Dynamic movements of ER in axonal growth cones of neurons transduced with**

**shDrebrin A+E.** Motoneurons expressing mCherry-ER were transduced with a shRNA lentivirus targeting both drebrin A and E and ER dynamic movements were imaged for 15 min at 2 sec intervals using an epifluorescence microscope. Related to Fig. 4.
